# Aortic valve disease augments vesicular microRNA-145-5p to regulate the calcification of valvular interstitial cells via cellular crosstalk

**DOI:** 10.1101/2022.11.29.518326

**Authors:** PR Goody, D Christmann, D Goody, D Nehl, K Becker, K Wilhelm-Jüngling, S Uchida, JB Moore, S Zimmer, F Bakhtiary, A Pfeifer, E Latz, G Nickenig, F Jansen, MR Hosen

## Abstract

**Rationale:** Aortic valve stenosis (AVS) is a major contributor to cardiovascular death in the elderly population worldwide. MicroRNAs (miRNAs) are highly dysregulated in patients with AVS undergoing surgical aortic valve replacement (SAVR). However, miRNA-dependent mechanisms regulating inflammation and calcification or miRNA-mediated cell-cell crossstalk during the pathogenesis of AVS are still poorly understood. Here, we explored the role of extracellular vesicles (EV)-associated *miR-145-5p*, which we showed to be highly upregulated upon valvular calcification in AVS in mice and humans.

**Methods:** Human TaqMan miRNA arrays identified dysregulated miRNAs in aortic valve tissue explants from AVS patients compared to non-calcified valvular tissue explants of patients undergoing SAVR. Echocardiographic parameters were measured in association with the quantification of dysregulated miRNAs in a murine AVS model. *In vitro* calcification experiments were performed to explore the effects of *EV-miR-145-5p* on calcification and crosstalk in valvular cells. To dissect molecular miRNA signatures and their effect on signaling pathways, integrated OMICS analyses were performed. RNA sequencing (RNA-seq), high-throughput transcription factor (TF) and proteome arrays showed that a number of genes, miRNAs, TFs, and proteins are crucial for calcification and apoptosis, which are involved in the pathogenesis of AVS.

**Results:** Among several miRNAs dysregulated in valve explants of AVS patients, *miR-145-5p* was the most highly gender-independently dysregulated miRNA (AUC, 0.780, p-value, 0.01). MiRNA arrays utilizing patient-derived- and murine aortic-stenosis samples demonstrated that the expression of *miR-145-5p* is significantly upregulated and correlates positively with cardiac function based on echocardiography. *In vitro* experiments confirmed that *miR-145-5p* is encapsulated into EVs and shuttled into valvular interstitial cells. Based on the integrated OMICs results, *miR-145-5p* interrelates with markers of inflammation, calcification, and apoptosis. *In vitro* calcification experiments demonstrated that *miR-145-5p* regulates the *ALPL* gene, a hallmark of calcification in vascular and valvular cells. EV-mediated shuttling of *miR-145-5p* suppressed the expression of *ZEB2*, a negative regulator of the *ALPL* gene, by binding to its 3’ untranslated region to inhibit its translation, thereby diminishing the calcification of target valvular interstitial cells.

**Conclusion:** Elevated levels of pro-calcific and pro-apoptotic EV-associated *miR-145-5p* contribute to the progression of AVS via the *ZEB2-ALPL* axis, which could potentially be therapeutically targeted to minimize the burden of AVS.

**Clinical Significance:** *What is known?:*

1. Aortic valve stenosis (AVS) is the most prevalent structural heart valve disease requiring surgical or interventional valve replacement. Currently, no medical treatment option is available to slow, halt, or reverse the progression of the disease.
2. AVS induces pressure overload on the left ventricle (LV), resulting in concentric hypertrophy and LV dysfunction.
3. AVS is not an exclusively degenerative disease that leads to fibrosis and calcification of the valve cusps but rather a chronic inflammatory disease, in which mechanical strain and shear stress lead to endothelial dysfunction and immune cell infiltration, which induces chronic inflammation, apoptosis and differentiation of valvular interstitial cells into osteoblast-like cells.
4. Increasing osteoblastic differentiation and the formation of macrocalcifications are hallmarks of the later stages of AVS.

*What is the new information we provide?:*

1. During aortic valve stenosis, expression pattern of vesicle-associated regulatory miRNAs is altered.
2. Patient-derived aortic valve tissue demonstrated an increased expression of *miR-145-5p* in humans, as well as in aortic valve explants from an experimental murine AVS model.
3. *MiR145-5p* contributes to calcification of the aortic valve through ZEB2, a transcriptional repressor of ALPL, in valvular interstitial cells.
4. Extracellular vesicular shuttling of *miR-145-5p* contributes to valvular cell-cell crosstalk and plays a role in the pathogenesis of AVS.

## Introduction

Aortic valve (AV) disease is a significant contributor to cardiovascular death worldwide and shows a prevalence of over 2% in cardiovascular patients over 60 years of age^1^. The 2-year mortality rate is greater than 50% when symptoms of severe aortic valve stenosis (AVS) are manifest, including dyspnoea, angina pectoris, or cardiac syncope^1–2^. While AVS is in part a degenerative disease, with “wear and tear” driving pathological fibrosis and calcification of the valve cusps, chronic inflammation is also a significant contributor to disease pathogenesis^1–2^. AVS can be divided into distinct phases. Initial endothelial damage, due to mechanical and shear stress, leads to infiltration of phospholipids (PL) and low-density lipoprotein particles (LDL). PL and LDL can be oxidized in the valve cusps, creating a pro-inflammatory milieu. Infiltration by monocytes and T-cells characterizes the initiation phase of the disease. Pro-inflammatory cytokines, secreted by classically activated macrophages and CD-8^+^ T-cells can induce apoptosis and differentiation of valvular interstitial cells (VIC) into osteoblast-like cells^3–4^. Arising cell debris acts as a further promotor of inflammation and can function as an initiator for microcalcification. Increasing osteoblastic differentiation and the formation of macrocalcification are indicative of the next phase of AVS, termed the propagation phase. Currently, no pharmacological treatments targeting either phase of the disease are available, and the only treatment option is surgical or interventional valve replacement (SAVR or TAVR)^5–7^.

MicroRNAs (miRNAs) are small noncoding RNAs that are involved in cardiovascular diseases^7–8^. When compared to healthy valves, miRNAs have been shown to be differentially expressed in aortic valve tissues from patients undergoing valve replacement surgery due to AVSRef. MiRNAs can also be packaged into extracellular vesicles (EV) to be transferred between different cells^8–10^. This phenomenon has been investigated in many diseases, including different types of cancer and in atherosclerosis^11–13^. We and others have demonstrated miRNA transfer between endothelial cells (EC), cardiomyocytes (CM), and smooth muscle cells (SMC), leading to direct genetic and phenotypic effects on target cells^8,14^. Recently, we showed that an increase in *EV-miR-122-5p* in patients with AVS represents a novel mechanism for the deterioration of cardiac function in patients following TAVR^8^. EVs harbor *miR-122-5p* and facilitate its shuttling into CM by direct interaction with a multifunctional RNA-binding protein (RBP), heterogeneous nuclear ribonucleoprotein U (hnRNPU), to regulate the viability of CM^8^. Furthermore, vesicular shuttling of *miR-30c-5p* is regulated by hnRNPU in a sequence-specific manner, which controls EC function and is augmented in CAD^14^. EVs have also been shown to play a role during AVS, with released EVs from VICs and macrophages acting as crystallization sites for microcalcification^15–18^. However, the horizontal transfer of vesicle-bound contents (e.g. miRNAs, proteins) and its effect on valvular calcification have not been well investigated.

The aim of our study was to explore whether tissue-resident miRNA content differ between patients with and without AVS. Further, we examined whether vesicular RNA contents are distinctive and how such differences impact disease progression through cell-cell communication via EVs. We demonstrated for the first time that vesicular *miR-145-5p* is upregulated in clinical and experimental settings of valvular calcification (i.e in AVS patients, our murine AVS model, and during *in vitro* calcification). Further functional and mechanistic studies revealed that *EV-miR-145-5p* is an important regulator of valvular calcification and osteoblastic differentiation of intestinal cells via vesicular shuttling.

## Methods

A detailed methods section is provided in the online Data Supplement.

### Study approval and Human specimen

All clinical samples and measurements were obtained after informed consent from patients following ethical approval by the ethics committee of the University of Bonn (approval number: AZ78/17). Aortic valve specimens were collected from patients undergoing SAVR for either severe aortic stenosis or aortic regurgitation in cooperation with the Department of Heart Surgery, University Hospital Bonn, Germany.

### Data availability

MiRNA array data are available from the Gene Expression Omnibus (GEO) under the accession number: GSE1905689. RNA-seq data are available from the GEO under the accession number GSE190539. The raw data of proteome and transcription factor arrays are provided as online data supplements. All further data that support the findings of this manuscript are available upon request from the corresponding author.

### Statistical analysis

Normally distributed continuous variables were presented as the mean ± standard deviation (SD). Continuous variables were tested for normal distribution using the Kolmogorov–Smirnov test. Categorical variables are given as frequencies and percentages. For continuous variables, the two-tail, unpaired Student t-test or Mann– Whitney U test were used for the comparison between the two groups. For the comparison of >2 groups, the one-way ANOVA with Bonferroni correction for multiple comparisons test was used. All tests were two-sided. Statistical significance was assumed when the null hypothesis could be rejected at p<0.05. Statistical analysis was performed with IBM SPSS Statistics version 20 (IBM Incorporation, USA) and GraphPad Prism 9 (GraphPad Inc, USA).

## Results

### Baseline characteristics and identification of differentially regulated miRNAs in valve tissue from AVS patients post-SAVR

To identify differentially expressed miRNAs in valve tissues, the AVS patients were characterized. Baseline characteristics indicated that comorbidities influencing the prognosis after SAVR [e.g., pulmonary disease and coronary artery disease (CAD)] were similar between the groups (AVS vs. AI). Of note, other cardiovascular risk factors such as type-II diabetes (Type II DM), body mass index (BMI), dyslipidemia, smoking history, as well as creatinine levels, were noted to be elevated in patients with AVS (Table 1). To identify differentially expressed miRNAs in valve tissue explants from AVS patients, we performed an unbiased RT-qPCR-based human miRNA array in the screening cohort (Figure 1A-B, Figure S1A). Explanted valves from patients with aortic insufficiency (AI, no AVS) due to dilatation of the aortic root or ascending aorta served as controls when valve cusps did not exhibit signs of calcification. Explanted valves from AVS patients displayed calcifications, increased collagen deposition, elastin, and fibrin in aortic valve tissue sections stained with alizarin red staining followed by light-microcopy imaging (Figure 1A). Interestingly, our array data revealed several miRNAs (*miR-145-5p, miR-let-7b, miR-1201, miR-145-3p, miR-29b, miR-126-3p, miR-29c, miR-126-5p, miR-133b, miR-518d, miR-127-5p*, and *miR-143-5p*) (Figure 1B-C) to be differentially expressed between AVS patients and controls after SAVR [Threshold values: |fold change| >1.5 and p-value <0.05 (FDR-adjusted)]. Of these differentially expressed miRNAs, *miRNA-145-5p* exhibited a more than 3-fold (p=0.001) increase in AVS tissues compared to those sourced from control patients (no-AVS). Said findings were recapitulated in a validation cohort via RT-qPCR, wherein miR*-145-5p* was significantly upregulated in AVS patients relative to controls (n=25 and n=10, respectively; p=0.01) (Figure 1D). The increased expression of *miR-145-5p* was independent of sex, with no significant difference being observed between male and female AVS patients in this validation cohort (Figure 1E).

**Table. 1.** Baseline characteristics of the study population

|  | **Control** | **AVS** |
| --- | --- | --- |
| Total population | 10 | 30 |
| **Clinical parameters** |  |  |
| Age (mean) | 68.6 | 69.7 |
| NYHA level | 3.3±0.5 | 3.1±0.5 |
| BMI | 24.3±6.7 | 25.2±3.7 |
| CAD | 5 (50) | 13 (43.3) |
| Hypertension | 8 (80) | 26 (86.6) |
| AVA in cm2 | 0.81 | >2.0 |
| Vmax in m/s | 4.45 | <1.0 |
| Pmean in mmHg | N.D. | 50.7 |
| LV function (mean) | 50.4 | 58.3 |
| eGFR (ml/min) | 54.2±15.8 | 51.6±20.0 |
| Mild-Severe mitral regurgitation | 7 (70) | 17 (70) |
| **Cardiovascular risk factors** |  |  |
| Type II DM | 0 (0) | 11 (36.7) |
| BMI (mean) | 24.2 | 29.288 |
| Dyslipidemia | 4 (40) | 23 (76.7) |
| Smoker | 4 (40) | 12 (40) |
| Creatininine lin mg/dl (mean) | 1.14 | 1.04 |

Baseline demographic, laboratory and echocardiographic parameters of the validation study population. P values reflect the comparison between two groups. NYHA, New York Heart Association; eGFR, estimated glomerular filtration rate; CAD, Coronary artery disease; Vmax, Velocity maximum, Pmean, mean pressure; BMI, Body mass index; AVA, Aortic valve area

**Figure 1.**
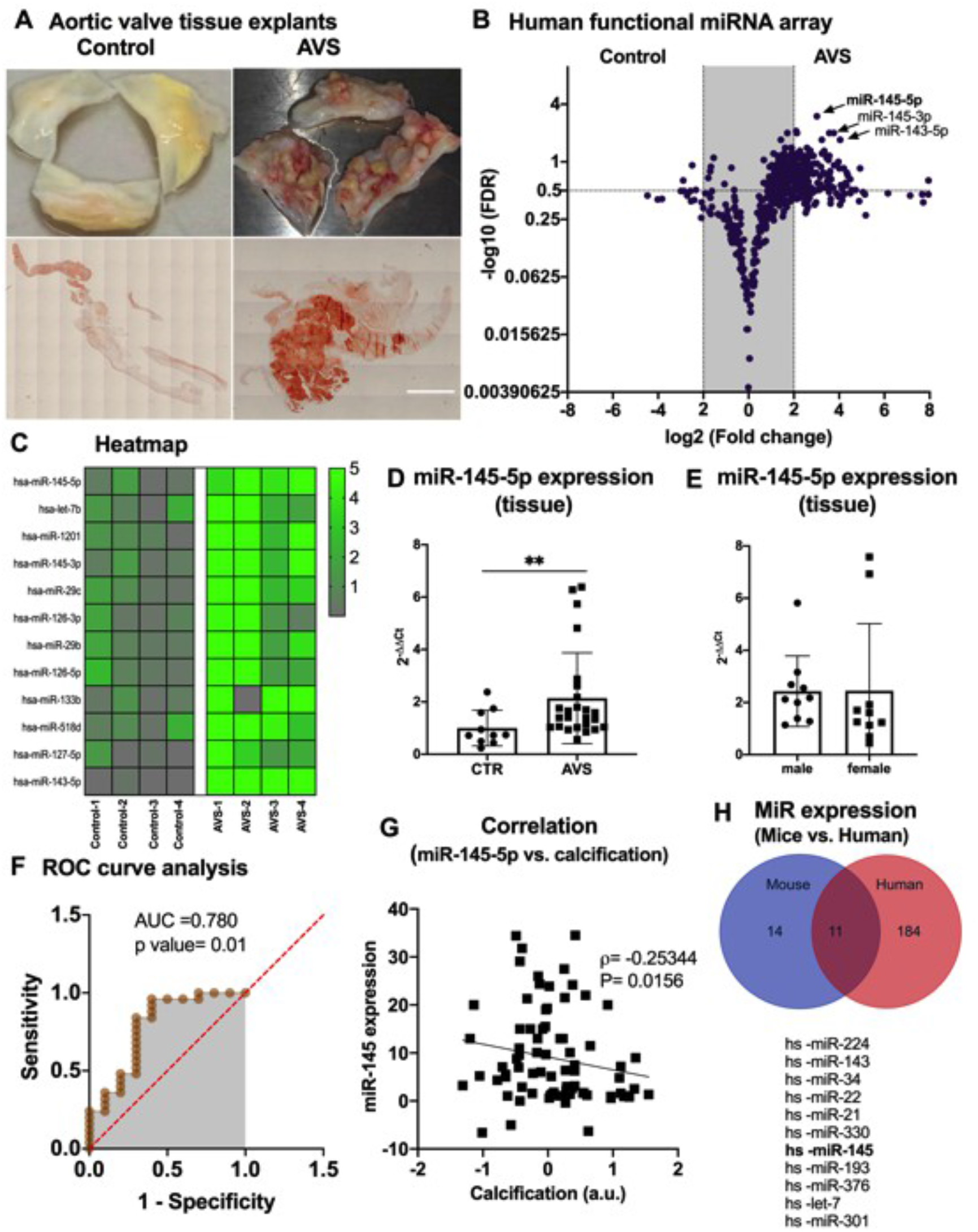
Investigation and profiling of miRNAs in stenotic aortic valves from patients who underwent surgical valve replacement with aortic valve stenosis. (A) Representative images of human calcified (AVS) and non-calcified (no AVS=AI) aortic valves explanted from patients undergoing aortic valve replacement surgery (upper part) and stained with alizarin red staining (lower part). (B) Volcano plot showing differentially regulated human miRNAs in explanted valve tissues derived from patients who underwent SAVR. Thresholds (black dotted lines) of a two-fold change and *p*-values (FDR-adjusted)<0.05 were used to distinguish the miRNAs of interest. n=4 for control (aortic insufficiency), n=4 for aortic valve disease (AVS). (C) Heatmap showing top-regulated miR expression of tissue-resident miRs were analyzed in explanted aortic valve tissues derived from controls (n=4) and AVS patients (n=4). (D) Expression of tissue-associated *miR-145-5p* was analyzed in control and AVS patients by qRT-PCRs. Data represent the mean ± SEM (\*\**p*<0.01, CTR, n=10, AVS n=30, by Student t-test, two-tail, unpaired). (E) Expression of tissue-associated *miR-145-5p* was analyzed in male and female patients by qRT-PCRs. Data represent the mean ± SEM (ns, *p*<0.06, Male, n=12, AVS n=12, by Student t-test, two-tail, unpaired). (F) ROC curve analysis for the prediction of expression of *miR-145-5p* in-patients with AVS and controls. (AUC, 0.780, **p-value 0.01). (G) Correlation analysis of *miR-145-5p* expression with the level of calcification. (Spearman coefficient of r=-0.25344, p=0.0156). (H) Venn diagram representing the dysregulated and common miRNAs in aortic valve samples from mice (sham, n=5, AVS, n=5) and patients with AVS (control, n=4, AVS, n=4). Mice were subjected to a wire injury to induce aortic valve stenosis *in vivo*. Aortic valves from 5 mice were pooled and analyzed via miRNA array, and 5 aortic valves from sham-operated mice were used as controls. *MiR-145-5p* was found to be amongst the upregulated miRNAs in both human and murine stenotic valves. The common miRNAs are shown in the list with a cut-off>2 fold, p-value<0.05 (FDR-adjusted). SAVR, surgical aortic valve replacement; AVS, aortic valve stenosis; miRNA, microRNA; ROC, receiver-operating characteristic curve.

Among the differentially expressed miRNAs, only *miR-145-5p* showed a significant diagnostic value in AVS patients when a ROC analysis was performed. This analysis suggested that *miR-145-5p* [AVS vs. controls] was a reliable predictor of the pathogenesis of disease (AUC=0.780, p=0.01; Figure 1F). We found that a significantly higher percentage of AVS patients demonstrated increased *miR-145-5p* expression. In addition, a correlation analysis of *miR-145-5p* expression with the level of calcification from AVS to controls suggests that *miR-145-5p* is a critical regulator of the calcification state of AVS patients (with a Spearman coefficient of r=-0.25344, p=0.0156) (Figure 1G).

### *MiR-145-5p* is highly upregulated in both human stenotic aortic valves and the murine AVS model

Most miRNAs are evolutionarily conserved, which helps in understanding their functions using model animals such as mice. To screen for species-conserved miRNAs in diseased aortic valve tissues, RT-qPCR-based miRNA arrays were performed using human (calcified tissue vs. control) and murine aortic valve tissue (wire injury vs. sham). Among eleven miRNAs that were shown to be differentially regulated (Figure 1H), *miR-145-5p* was expressed highly in both humans and mice. Based on the above-mentioned data, we sought to investigate the functional importance and molecular mechanism of *miR-145-5p* in AVS.

To further investigate the role of *miR-145-5p* in AVS, we employed a graded wire-injury model of AVS in mice (Figure 2A). A wire-induced aortic valve injury led to the development of severe stenosis, as demonstrated by elevated blood flow velocities four weeks after the operation (Figure 2A-B). The ejection fraction (EF) and cardiac output were inversely regulated in these mice, suggesting that there is an AVS-induced decrease of EF (Figure 2C-D) in comparison to baseline (when compared to sham-operated mice). RT-qPCR analysis revealed that *miR-145-5p* was highly upregulated in mice with AVS (Figure 2E).

**Figure 2.**
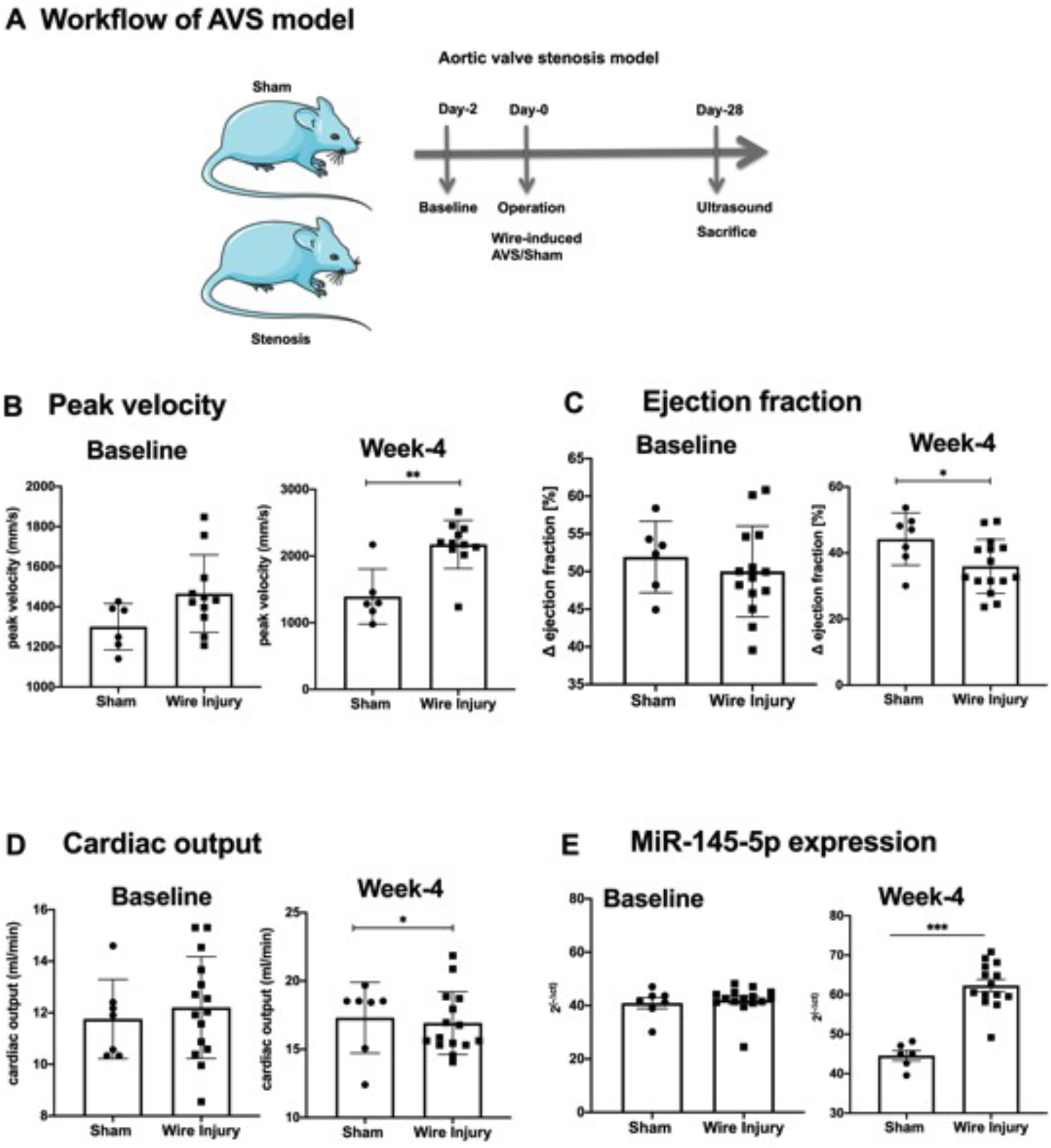
*MiR-145-5p* is differentially regulated in a murine model of aortic valve stenosis. (A) Timeline of the operation used to induce aortic valve stenosis in a murine model. The induction of aortic valve stenosis in mice is achieved by inserting a coronary wire into the left ventricle and rotating it. (B) Peak velocities over the aortic valve after sham operation or wire injury show induction of stenosis, as visualized with peak velocity, 4 weeks after surgery. Statistical significance was shown between, Sham-day-1 vs. Sham-week-4 and Stenosis-day-1 vs. Stenosis-week-4 (**p<0.01, sham, n= 13; stenosis, n= 13; SEM; One-way ANOVA with Bonferroni multiple comparisons test). (C) Change in the ejection fraction after sham operation or operation creating severe stenosis on day-1 and week-4, as compared to before the surgery. Statistical significance was shown between, Sham-day-1 vs. Sham-week-4 and Stenosis-day-1 vs. Stenosis-week-4 (*p<0.05, sham, n=13; stenosis, n=13; SEM; One-way ANOVA with Bonferroni multiple comparisons test). (D) Cardiac output after sham operation or operation creating severe stenosis on day 1 and week 4 (*p<0.05, sham, n=13; stenosis, n=13; SEM; One-way ANOVA with Bonferroni multiple comparisons test). (H) *MiR-145-5p* expression, as determined by RT-qPCR, in valvular tissue after sham operation or operation creating severe stenosis on day 1 and week-4 (*p<0.05, sham, n=13; stenosis, n=13; SEM; One-way ANOVA with Bonferroni multiple comparisons test). AVS, aortic valve stenosis.

### *MiR-145-5p* expression in interstitial cells is higher than endothelial cells in valve tissues

To elucidate the mechanism of action of *miR-145-5p*, we established several *in vitro* culture models. We established an improved isolation protocol for valvular endothelial cells (VECs) and valvular interstitial cells (VICs) from human calcified and non-calcified AV tissues explanted during SAVR (Figure 3A). In brief, human VICs and VECs (patVIC and patVEC) were isolated from explanted AVs using multiple steps of collagenase digestion and CD105 (Endoglin) Magnetic Activated Cell Sorting (MACS) for EC purity (Figure 2B). We further obtained human VICs and VECs (hVIC and hVEC) from a healthy young donor who died from a non-cardiovascular-related event. Patient-derived and commercially availabe cells were characterized via the expression of different endothelial and interstitial cell markers via qRT-PCR. A comparison of characteristic marker expression in these cells was performed (Figure S2A-D) to confirm cellular identity of isolated and commercially available valve cells. Immunofluorescence staining for prototypical endothelial and interstitial cell markers and gene expression were performed by qRT-PCR (Figure 3B-E, Figure S2A-D). PatVICs and hVICs stained positively for α-SMA, which was also further confirmed by qRT-PCR of these cells (Figure 3B-E). In contrast, patVECs and hVECs showed high expression levels of endothelial markers, such as vWF, PECAM1, and CDH5 (Figure S2A-D), thus further confirming that our MACS-based isolation was valid and reproducible.

**Figure 3.**
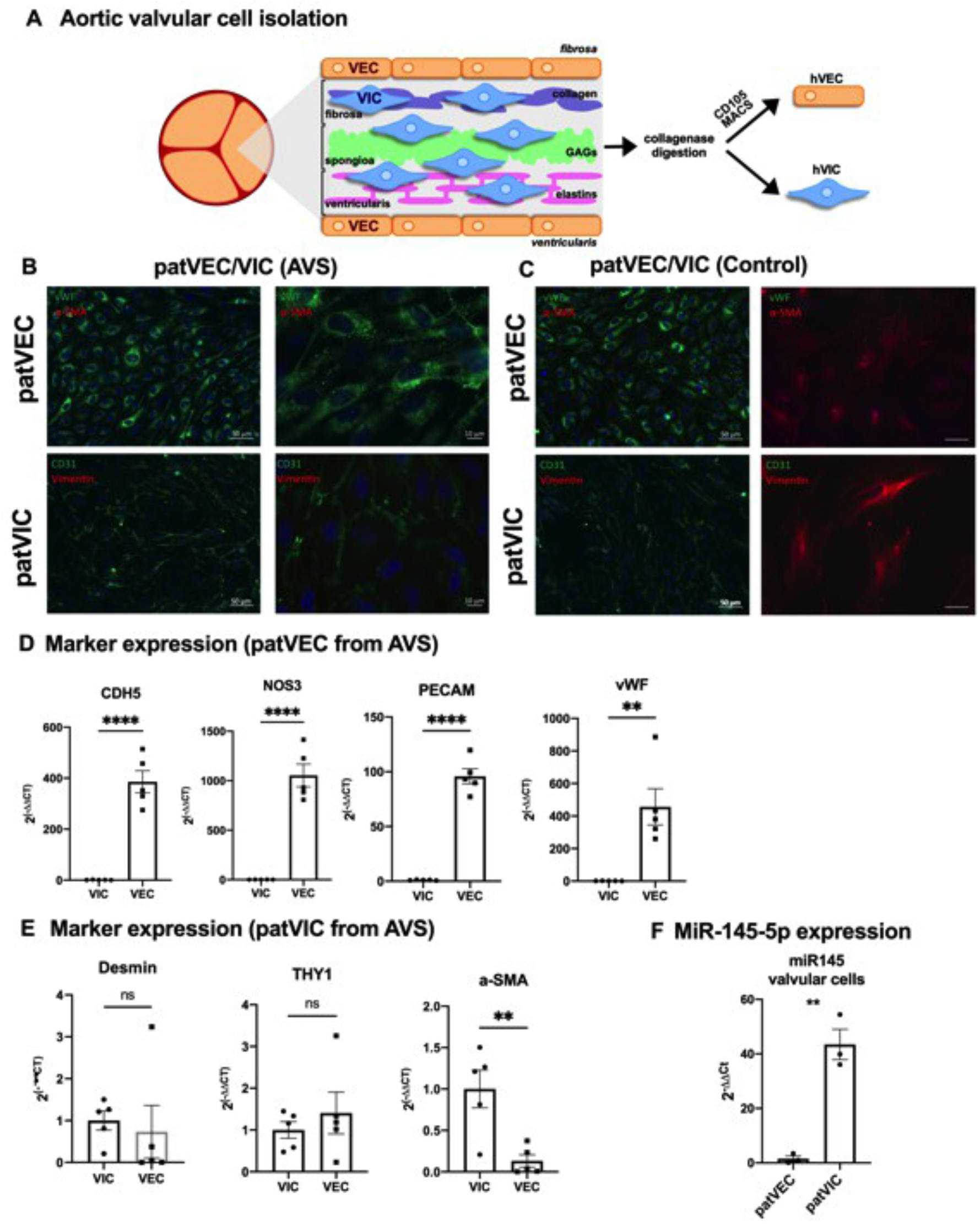
Isolation and characterization of patient-derived valvular cells. (A) Schematic representation of the structure of valvular matrix and isolation process of valvular cells from aortic valve tissue explant after SAVR. Minced aortic valve tissues have been subjected to collagenase digestions overnight at optimal cell culture conditions under gentle rotations. MACS-affinity-based selection was performed to isolate VECs, whereas; a preplatig step is required for VICs to separate from the remaining cell mixture. (B-C) Characterization of isolated aortic valve cells by using different surface markers, corresponding to VECs-and VICs-derived from AVS patients that underwent SAVR and compared with the corresponding controls. Representative immunofluorescence images of isolated VECs (patVECs) and VICs (patVICs) by using their characteristic markers. (D-E) Expression of characteristic markers of isolated valvular cells, (VECs, and VICs), isolated from patients with AVS post SAVR. Data are depicted as Student t-tests. (****p<0.0001, **p<0.01, n=5, two-tailed, unpaired). Expression of *miR-145-5p* was normalized to the internal control gene, *RNU6*. (F) Expression of *miR-145-5p* was measured in VECs and VICs isolated from patients with AVS after SAVR. Data are depicted as Student t-test. (**p <0.01, n=3, two-tailed, unpaired). Expression of *miR-145-5p* was normalized to the internal control gene, *RNU6*. VICs, valvular interstitial cells; VECs, valvular endothelial cells; SAVR, surgical valve replacement; vWF, von-Willebrand factor; a-SMA, alpha-smooth muscle actin; DAPI, 4’,6-Diamidin-2-phenylindol; PECAM1, platelet endothelial cell adhesion molecule; CDH5, cadherin 5/VE-Cadherin; NOS3, nitric oxide synthase 3/eNOS; DES, desmin; THY1, Thy-1 Cell Surface Antigen/CD90.

To investigate the cell-specific expression of *miR-145-5p* isolated from AV-tissues after SAVR, we quantified *miR-145-5p* expression in VECs and VICs. Our qRT-PCR quantification revealed that VICs express a significantly higher level of *miR-145-5p* in comparison to VECs (Figure 3F) suggesting that *miR-145-5p* may play a role in the pathogenesis of AVS in mice and humans through its expression in specific valve cells.

### AVS increases the level of *miR-145-5p* in aortic valve tissue and EVs derived from plasma of patients

Recent studies have shown that different extracellular vesicle (EV) populations (e.g. exosomes or small EVs, large EVs, apoptotic bodies) act as vehicles to transfer short or long RNAs into nearby or distant recipient cells and play a role in cellular crosstalk^8,14^. Encapsulation of miRNAs into small or large EVs provide dramatic resistance to blood- or tissue-resident exonucleases. MiRNAs can also be secreted bound to LDLs (low-density lipoproteins, HDL (high-density lipoproteins), and RBPs (e.g., argonaute, hnRNPU, hnRNPUA2B1, hnRNPK, HuR)^14–15^.

The characterization of EVs derived from AV tissue or plasma was performed according to the current guidelines of the International Society of Extracellular Vesicles (ISEV) using nanoparticle tracking analysis (NTA), transmission electron microscopy (TEM), and immunoblotting for vesicular markers, including tetraspanins (Flotillin, CD63, CD81) (Figure 4B-D).

**Figure 4.**
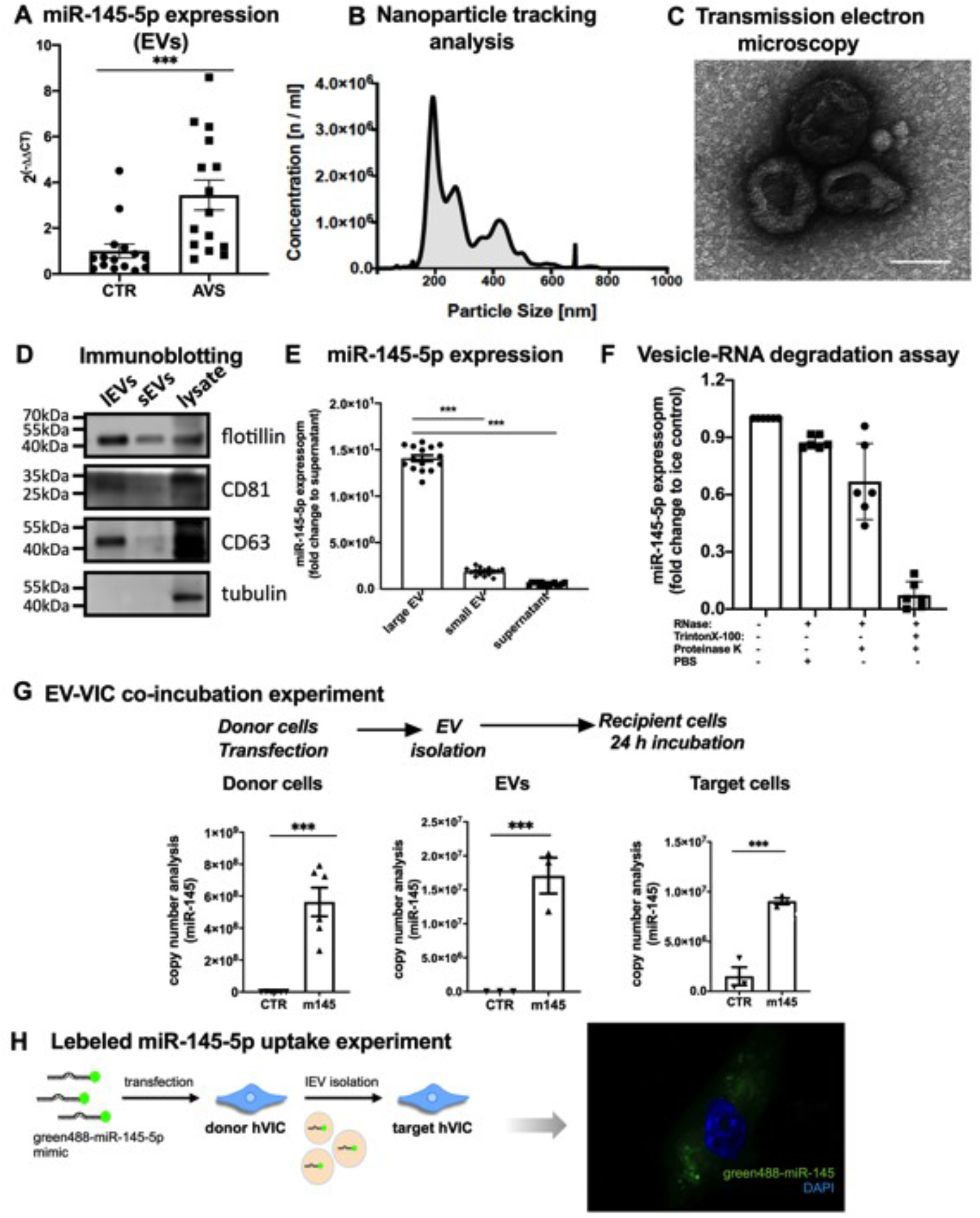
*MiR-145-5p* may regulate the calcification of recipient VICs via vesicular transfer. (A) Expression of *miR-145-5p* was measured in large EVs isolated from aortic valve tissues derived from patients with AVS and corresponding controls. Taqman-based expression analysis was performed to determine *miR-145-5p* in large EVs. Data is depicted with Student t-tests. (***p<0.001, n=15, two-tailed, unpaired). Expression of *miR-145-5p* was normalized to the internal control gene, *RNU6*. Large EVs were isolated by using centrifugation at 20,000 g, according to the protocol published by our group and others (20-21, 29-31). (B) NTA was used to determine the diameter (~200–700 nm) and concentration of large EVs isolated from valve tissue from patients undergoing SAVR (C) TEM image (85,000 x magnification) of pelleted large EVs (diameter ~300–700 nm) derived from the valve tissue of patients with AVS. (D) Western blot analysis of the expression of small and large EV-markers. (E) *MiR-145-5p* expression was assessed in different EV populations isolated from VIC cultures by qRT-PCR (\*\*\**p*<0.001, n=15, by 1-way ANOVA with Bonferroni correction for multiple comparisons test). Cel-miR-39 was used for normalization. EVs and exosomes were isolated by centrifugation at 20,000 *g* and 100,000 *g*. (F) Vesicle-RNA degradation assays. VIC-derived EVs were treated in parallel using different conditions followed by RNase A digestion. *MiR-145-5p* was quantified by qRT-PCR (*p<0.05, compared with the untreated group; ns: not significant, n=3, by 1-way ANOVA with Bonferroni correction for multiple comparisons test). (G) Co-incubation experiments with large EV isolated from miR-145 overexpressed VIC and control. VICs were transfected with control, *miR-145-5p* mimic and large EVs were isolated and co-incubated with VICs under similar treatment conditions to quantify the differential *miR-145-5p* transfer via large EVs to the target cells. *MiR-145-5p* expression was assessed in donor VICs, isolated large EVs, and target VICs that were treated with large EVs, using copynumber analysis (***p<0.001, n=6, by 1-way ANOVA with Bonferroni multiple comparisons test). (H) EV-incorporated *miR-145-5p* uptake experiments in VICs. PKH26-labeled large EV and green fluorescent 488-labeled *miR-145-5p* were coincubated for 24 hours to allow incorporation of EV-associated *miR145-5p* into recipient VICs. EVs, extracellular vesicles; AVS, aortic valve stenosis; TEM, transmission electron microscopy; NTA, nanoparticle-tracking analysis; VECs, valvular endothelial cells; VICs, valvular interstitial cells.

To identify the mode of transport of *miR-145-5p*, we determined the plasma component of AVS-patients in which *miR-145-5p* may be detected. After isolation of large EVs, small vesicles (also known as exosomes), and vesicle-free plasma (Figure S3A)^8, 14, 25–26^, the large EV population (170-800 nm) isolated from aortic valve tissues explanted from patients with AVS after SAVR demonstrated a significantly higher level of *miR-145-5p* expression in comparison to control (vesicle-free plasma) (Figure 4A), suggesting that *miR-145-5p* is contained in large EVs. To verify *miR-145-5p* secretion and transport in large EVs, a vesicle-RNA degradation assay was performed (Figure 4E-F). We found that proteinase K digestion before treatment with RNase did not affect *miR-145-5p* levels. In contrast, treatment with Triton X-100, which acts as a detergent to disrupt the phospholipid membrane of vesicles, before treatment with RNase led to near-complete degradation of *miR-145-5p*. These findings indicate that extracellular *miR-145-5p* may be incorporated into large EVs, protecting the RNA from resident or circulating nucleases. Altogether, these data suggest that extracellular *miR-145-5p* is predominantly encapsulated and secreted in large EVs.

### Vesicular shuttling augments the expression level of *miR-145-5p* in recipient valvular interstitial cells

As dysfunction of VICs is crucial to the development of AVS in the murine AVS model and human, we explored whether VICs can transfer *miR-145-5p* into target VICs (other neighboring VICs) and established a co-culture model of donor and recipient VICs to investigate intercellular communication via EVs. After transfecting VICs with *miR-145-5p* mimic, large EVs were isolated and incubated with target VICs, revealing that *miR-145-5p* is upregulated in target VICs in both control and mimic-transfected cells (Figure 4G). This result suggests that horizontal EV-mediated cell-to-cell transfer of *miR-145-5p* increases the levels of *miR-145-5p* in target VICs.

To examine whether the intercellular transfer of *miR-145-5p* occurs via shuttling by large EVs in a paracrine manner, fluorescently labeled large EVs were isolated from VICs and incubated with acceptor VICs. The results show that the fluorescently labeled EVs can be internalized into recipient VICs (Figure S3B). Furthermore, large EVs isolated from donor VICs transfected with fluorescently labeled *miR-145-5p* confirmed the uptake of EV-*miR-145-5p* into recipient VICs (Figure 4H), indicating that EV-mediated shuttling of *miR-145-5p* could act as an intercellular mediator of calcification within the aortic valve during stenosis progression.

### *MiR-145-5p* regulates pro-calcification marker genes in VICs

Recent studies have reported that miRNAs can influence the calcification process in vascular and valvular cells via regulation of pro-calcific genes or proteins^31–32^. To examine whether the calcification of VICs is mediated by *miR-145-5p*, we performed *in vitro* calcification experiments. The level of calcification was assessed by utilizing alizarin red staining to quantify matrix calcium on deposition (Figure 5A). To further elucidate any direct involvement of *miR-145-5p* during calcification of VICs, we proceeded to assess the expression of calcification marker genes via qRT-PCR in cells treated with either a *miR-145-5p* mimic or inhibitor (Figure 5B, Figure S4A-B). Interestingly, upon silencing of *miR-145-5p*, downregulation of calcific genes, including alkaline phosphatase, biomineralization associated (*ALPL*), zinc finger E-box binding homeobox 2 (*ZEB2*), RUNX family transcription factor 2 (*RUNX2*), secreted phosphoprotein 1 (*SPP1*), matrix Gla protein (*MGP*), KLF transcription factor 4 (*KLF4*), and SMAD Family Member 5 (*SMAD5*), was observed,. These data suggest that *miR-145-5p* could regulate key genes in calcification processes that can be partly abrogated by overexpression of *miR-145-5p* (Figure S4A-B).

**Figure 5.**
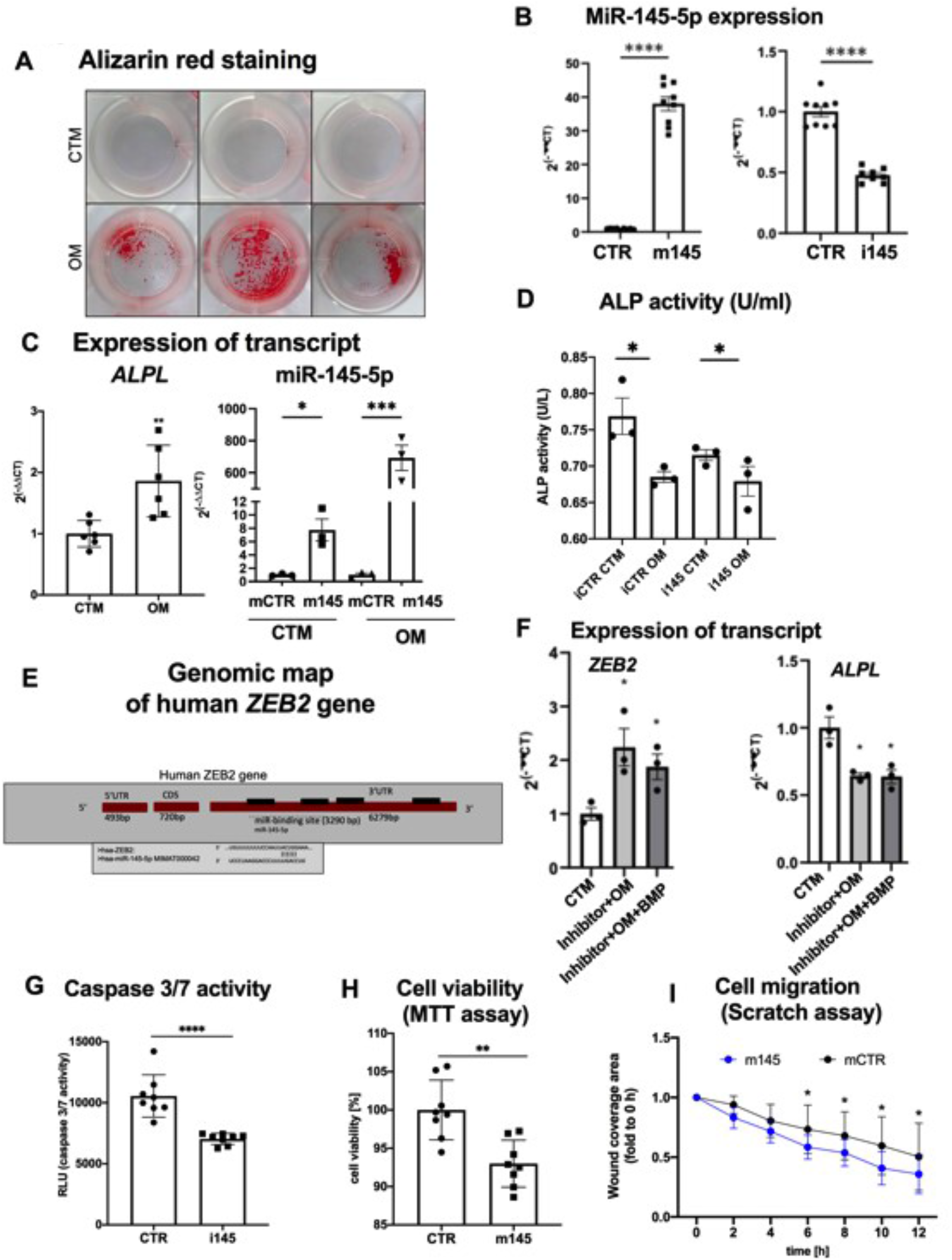
*MiR-145-5p* is a crucial regulator of valvular calcification and cellular function. (A) *In vitro* calcification experiment in VICs followed by quantification via Alizarin red staining on day 7. The deposition of calcium in the cellular matrix was stained and quantified with alizarin red after induction of *in vitro* calcification in VICs purchased from a commercial vendor (representative images of n=3). (B) Overexpression and knockdown quantification of *miR-145-5p* by small oligonucleotides in VICs. Expression data for *miR-145-5p* in VICs have been compared with control and depicted as ****p<0.001, n=9, by 1-way ANOVA with Bonferroni multiple comparisons test. (C) Taqman-based quantification of gene expression of ALPL as a hallmark of calcification of VICs and *miR-145-5p* after induction of *in vitro* calcification by using osteogenic medium (OM) for 7 days. Expression data for ALPL mRNA in VICs and *miR-145-5p* has been compared with control medium and control *miR-145-5p* mimic and depicted as*p<0.05, **p<0.01, ***p<0.001, n=3-6, by 1-way ANOVA with Bonferroni multiple comparisons test. (D) Quantification of ALP activity in VICs after transient *miR-145-5p* overexpression or inhibition and simultaneous incubation with OM (n=3, 2-way ANOVA with Bonferroni multiple comparisons test). (E) Genomic map of the ZEB2 gene, a known regulator of ALPL, with predicted binding sites for *hsa-miR-145-5p*, in the 3’-UTR. ZEB2 is a known repressor of ALPL and contains multiple binding sites for numerous miRs, including, miR-145-5p. (F) Expression of ZEB2 and ALPL mRNA in VICs after induction of *in vitro* calcification for 7 days. Data is depicted as *p<0.05, n=3, by 1-way ANOVA with Bonferroni multiple comparisons test. (G) The level of apoptosis was assessed via caspase 3/7 activity after incubation with H2O2 (100 μM) for 24h. The data was analyzed by 1-way ANOVA with Bonferroni’s multiple comparison test. (****p<0.001, n=8). (H) Cell viability was determined by using MTT assays (incubation with H2O2 (100 μM)). Data was analyzed by 1-way ANOVA with Bonferroni’s multiple comparison test. (**p<0.01, n=8). (I) Cell migration via scratch-wound was determined by using bright-field microscopy after overexpression of miR-145-5p and control at different time points. Cell-free areas were measured to assess the migration capacity of VICs at 0h, 2h, 4h, 6h, 8h, 10h, and 12h time points after scratch-wound. The data was analyzed by 1-way ANOVA with Bonferroni’s multiple comparison test. (*p<0.05, n=3). ZEB2, Zinc Finger E-Box Binding Homeobox 2; ALPL, Alkaline Phosphatase; CTM, control media, OM, osteogenic medium; UTR, untranslated region; VICs, valvular interstitial cells; miRs, microRNAs; H2O2, hydrogen peroxide; AVS, aortic valve stenosis.

Among the dysregulated genes involved in calcification processes, *ALPL* was observed to be upregulated upon induction of calcification in *miR-145-5p* overexpressed VICs (Figure S4B-C). The inverse correlation between *miR-145-5p* and *ALPL* was further confirmed when VICs were incubated with osteogenic medium (OM) to induce calcification (Figure 5C), indicating that *miR-145-5p* regulates the *ALPL* expression in a dose-dependent manner (Figure S4C). To examine whether *miR-145-5p* regulates the transcription of the *ALPL* or ALP activity itself, we quantified alkaline phosphatase activity upon transfection of VICs with mimic and inhibitors followed by induction of calcification *in vitro*. Upon knockdown of *miR-145-5p* in VICs, we observed reduced enzymatic activity of ALP (Figure 5D), suggesting that *miR-145-5p* may modulate the expression of *ALPL*, thus diminishing ALP activity.

### *MiR-145-5p* may regulate calcification of VICs through ZEB2 and tissue nonspecific alkaline phosphatase (*ALPL*)

Several studies revealed that *ALPL* transcription can be regulated by numerous transcription factors (TFs). For example, ZEB2 is a known repressor of the *ALPL* and was reported to be increased in healthy heart tissues, including AV tissue^30–32^. Given that *miR-145-5p* was reported to bind to *ZEB2* to regulate cellular gene expression during epithelial-mesenchymal transition (EMT) in metastasis^42^, we sought to further characterize its regulation by *miR-145-5p*. To study whether *ZEB2* has binding sites for *miR-145-5p*, we analyzed the genomic architecture of the *ZEB2*. Interestingly, genomic analysis and target prediction of *miR-145-5p* binding demonstrated that the *ZEB2* contains potential binding sites for numerous miRNAs, including *miR-145-5p*, 3290 bp upstream of the first two exons in its 3’-UTR (untranslated region). This suggests that *miR-145-5p* may bind to *ZEB2* (Figure 5E), consequently regulating the expression of its cognate *ALPL* mRNA and thus protein synthesis.

Given that ALPL is a central player in calcification^29–32^, we sought to further characterize its regulation by *miR-145-5p*. We reciprocally quantified the expression of *ALPL* and its transcriptional repressor*, ZEB2*, in the same experiments upon knockdown of *miR-145-5p*. The induction of calcification by osteogenic medium showed that the expressions of *ZEB2* and *ALPL* are inversely correlated, an effect also seen upon induction of *ZEB2* by BMP (bone morphogenic proteins) (Figure 5F), suggesting that *miR-145-5p* regulates the transcription of the *ALPL* gene via its transcriptional repressor *ZEB2* by binding to the 3’-UTR of *ZEB2*. To examine whether this regulation has any effect on the cellular function of VICs, we performed experiments focusing on cellular viability and functions; namely, the activation of caspase 3/7 for apoptosis, cell viability via MTT reduction, and cell migration via scratch-wound healing assays upon silencing or overexpressing *miR-145-5p*. The results revealed that *miR-145-5p* can regulate apoptosis and viability of VICs as a pro-apoptotic miRNA (Figure 5G-I). Taken together, our data suggest that *miR-145-5p* exerts a pro-apoptotic effect that is a prerequisite of the initiation of valvular calcification in AVS.

### RNA sequencing reveals that *in vitro* calcification alters calcific gene and miRNA expression profiles in valvular interstitial cells

To assess calcification-induced gene expression signatures in the VICs, we performed bulk RNA sequencing of calcified VICs upon incubation with OM for 7 days and controls (without OM), resulting in a robust differential expression of pro-calcific markers, e.g., *ADAMS18, RUNX2, ALPL, COL11A1*, and *MMP13* (Figure 6A). We further analyzed the miRNA expression profile of these VICs and found *miR-145-5p* among the dysregulated miRs (Figure 6B), providing further confirmation of our *in vitro* and *in vivo* data, which identify *miR-145-5p* to be involved in the calcification process of VICs. Further bioinformatics analysis via DAVID (Database for Annotation, Visualization and Integrated Discovery) and KEGG (Kyoto Encyclopedia of Genes and Genomes) revealed that induced pathways involve osteogenesis and apoptosis among the top-regulated pathways, providing further evidence of the calcification potential of VICs (Figure S5A).

**Figure 6.**
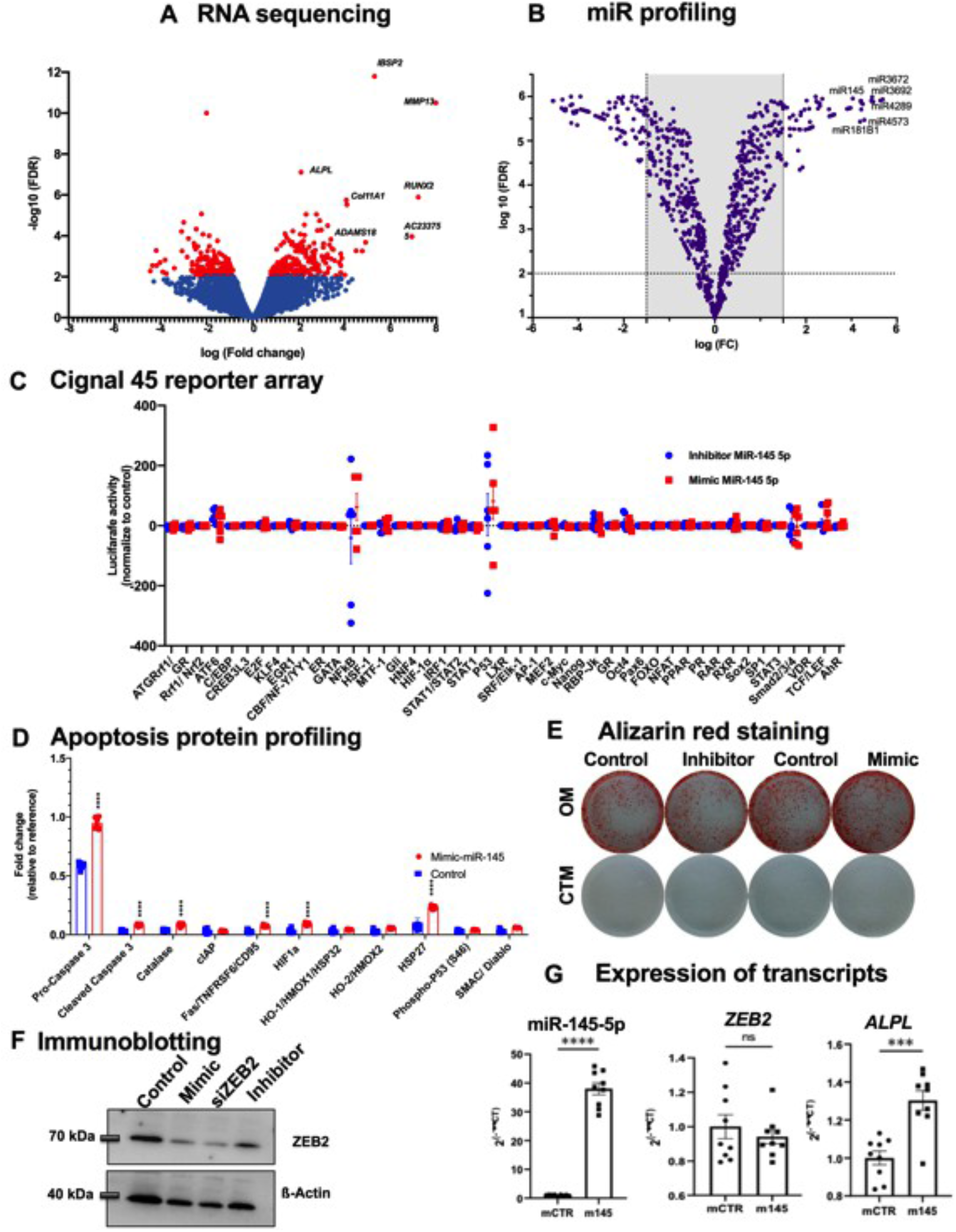
Transcriptomic profiling of calcified VICs confirms that miR-145-5p regulates calcification and apoptosis by regulating *ALPL2* and other calcific genes. (A) Volcano plot of RNA sequencing data of calcified human VICs (7 days) vs. controls that demonstrated an upregulation of genes that are central to calcification, such as *ALPL, RUNX2, MMP13, ADAMTS18, COL11A1*, and *AC233755*, etc. Furthermore, an upregulation of known inflammation-, and apoptosis-regulated genes upon induction of calcification for 7 days was observed (n=4; SEM; FDR-adjusted). (B) Volcano plot with profling of dysregulated miRNAs in calcified VICs (7 days) in comparion to control VICs with a cut off 1.5 fold (n=4; SEM; FDR-adjusted). The top and statisitcally significantly upregulated miRNAs are represented. (C) A high throughput transcription factor activity reporter luciferase assay upon oligoneucleotide-mediated depletion or overexpression of miR-145 vs. control in VICs was performed 48h after transfection. Depicted are the activities of 45 transcription factors in inibitor, *miR-145-5p* treated cells and control cells (control-treated) 48h after transfection with the reporter constructs. Pro-Caspase-3, Caspase-3, HSP27, HIF1a, and other apoptosis-related pathways are significantly upregulated. Data represent mean ± SEM (****p<0.0001, n≥6). (E) Representative bright-field images of *in vitro* calcification experiment in VICs followed by quantification via Alizarin red staining on day 7. The deposition of calcium in the cellular matrix quantified and stained with alizarin red after induction of *in vitro* calcification in VICs purchased from a commercial vendor (representative images of n=3). (F) Western blotting for ZEB2 protein after transfection with control, *miR-145-5p* inhibitor (Inhibitor), siRNA against ZEB2 (siZEB2) and *miR-145-5p* mimic (Mimic). (G) Overexpression data of *miR-145-5p, ZEB2* mRNA, and *ALPL* in VICs upon transfection and subsequent inducton of calcification for 7 days to assess the expression of the transcripts (^ns^p>0.05, ***p<0.001, ****p<0.0001, n=9, by 1-way ANOVA with Bonferroni multiple comparisons test). ZEB2, Zinc Finger E-Box Binding Homeobox 2; RUNX2, Runt-related transcription factor 2; ALPL, Alkaline Phosphatase; HSP27, Heat shock protein 27; HIF1a, Hypoxia-inducible factor 1-alpha; CTM, control media, OM, osteogenic medium; UTR, untranslated region; VICs, valvular interstitial cells; miRs, microRNAs; AVS, aortic valve stenosis.

### *MiR-145-5p* may exert its function via the *miR-145-5p-ZEB2-ALPL* axis in calcific valvular interstitial cells

To gain further insight into cellular pathways related to calcification regulated by *miR-145-5p*, an unbiased high throughput pathway reporter array for transcription factor activitiy was performed. VICs were either transfected with mimic or inhibitor of *miR145-5p* and control. The cells were then transfected with two plasmids, one carrying a firefly luciferase downstream of a transcription factor binding site and a renilla luciferase under the control of a CMV promoter as transfection control. After 24 h, transcription factor activity was calculated by luminescence measurements and normalized with control (Figure 6C, Figure S5B). Among the highly active transcription factors, *NFkB (Nuclear Factor Kappa B Subunit 1), STAT1 (Signal Transducer And Activator Of Transcription 1), STAT3 (Signal Transducer And Activator Of Transcription 3), P53 (cellular tumor antigen p53*) were dysregulated in the reporter array analysis (Figure 6C, Figure S5B, Table S1).

To examine how *miR-145-5p* regulates apoptosis in a more unbiased manner, we performed a proteome profiler assay with antibodies against 35 apoptosis-related proteins with VIC lysates after knockdown of *miR-145-5p* and or control (Figure 6D, Figure S5C, Table S2). Interestingly, several pro-apoptotic proteins (pro-caspase-3, cleaved caspase-3, fas/TNFRSF6, p53, etc.) were increased, whereas antiapoptotic proteins (hsp27, catalase, HIF-1α, etc.) were decreased. The dysregulation of these apoptotic proteins upon *miR-145-5p* knockdown suggests that miR-145-5p regulates viability and apoptosis of VICs during calcification (Figure 6D). These data are in line with our pathaway analysis and reporter arrays, as these proteins are known to modulate different stages of calcification. Finally, we assessed whether overexpression or knockdown of *miR-145-5p* supports our data and whether overexpression of *miR-145-5p* rescues the pro-apoptotic/calcific effect of ZEB2 and ALPL. We repeated calcification in VICs upon overexpression and knockdown of *miR-145-5p* followed by alizarin red staining and found that overexpression of *miR-145-5p* directly increases calcification (Figure 6E). As expected, when the expression of ZEB2 protein was quantified via immunoblotting with *miR-145-5p* mimic and siRNA against ZEB2, we found reduced expression of ZEB2 in mimic- and siZEB2-treated VICs, whereas the expression was increased in inhibitor-treated VICs, suggesting that *miR-145-5p* directly regulates ZEB2, which acts on *ALPL* expression as a transcriptional repressor (Figure 6F). To determine the involvement of miR-145-5p-ZEB2-ALPL as a mediator of the calcification process in VICs, we further quantified mRNA expression, which supports our finding that overexpression of *miR-145-5p* suppresses ZEB2, augmenting the expression of *ALPL* in VICs (Figure 6G).

Taken together, the above-mentioned data demonstrate that regulation of ZEB2-ALPL by *miR-145-5p* triggers apoptosis and calcification of VICs and that the level of expression of *miR-145-5p* is important for this mechanism, which acts in a *miR-145-5p-ZEB2*-ALPL-dependent manner (Figure 7). Overexpression of *miR-145-5p* in the AV induces inflammation, calcification, and apoptosis leading to stiffening of the aortic valve cusps and narrowing of the aortic valve orifice, ultimately inducing pressure overload of the LV and valvular heart failure.

**Figure 7.**
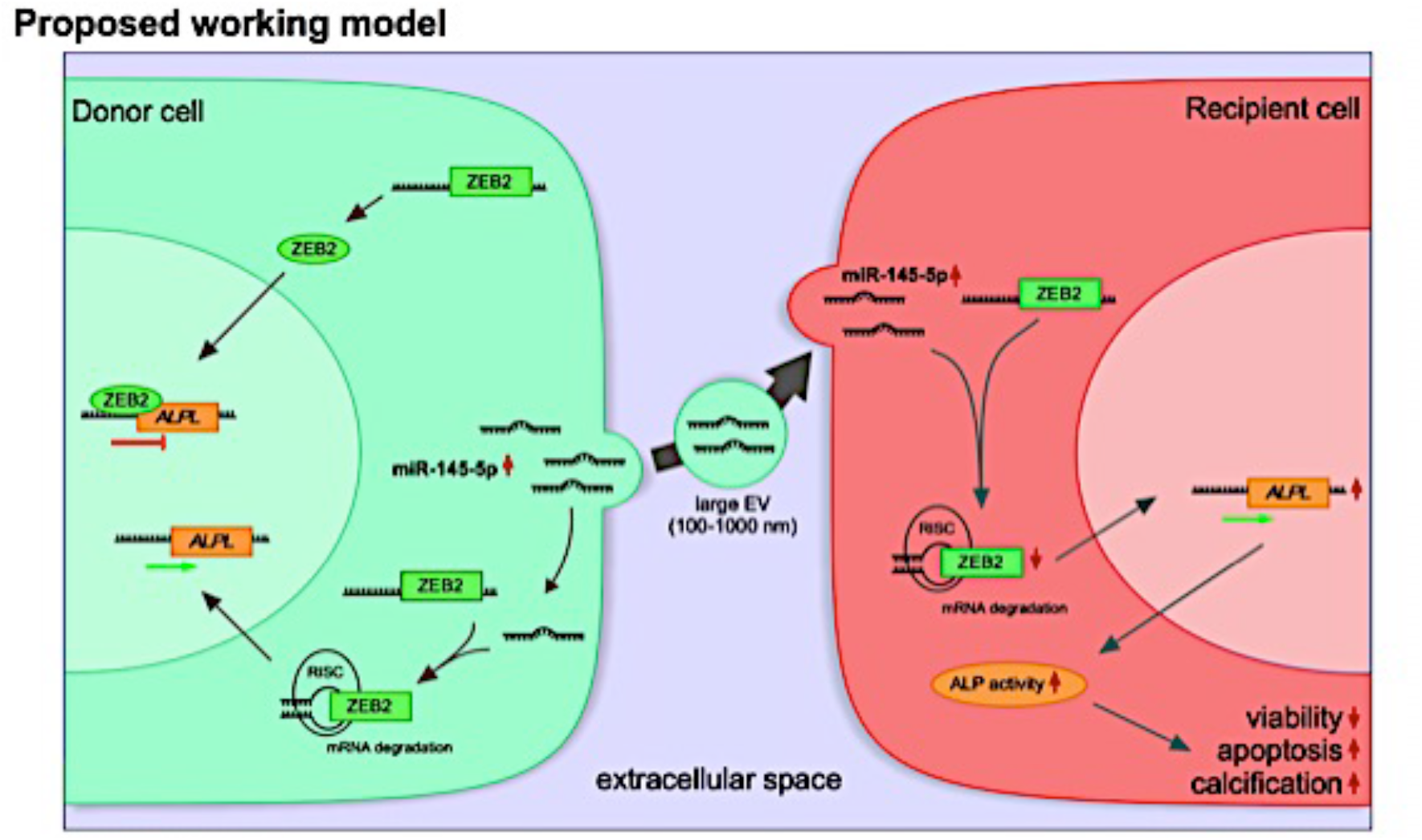
Large extracellular vesicular *miR-145-5p* regulates calcification and apoptosis of valvular interstitial cells by regulating *BCL2* and other apoptotic genes. The model proposed for *miR-145-5p* action, in which it is shuttled to the recipient cells via *ZEB2-miR-145-5p* conjugates to regulate calcific gene networks by binding to 3’-UTR, such as ALPL, to control the inflammation, calcification, and apoptosis of VICs. ICs, valvular interstitial cells; miRs, microRNAs; RISK, RNA-induced silencing complex; UTR, untranslated region; ZEB2, Zinc Finger E-Box Binding Homeobox 2; ALPL, Alkaline Phosphatase.

## Discussion

The data from the present study demonstrate a newly identified role for EV-miRNAs that are differentially expressed and associated with calcification in AVS patients. *Mir-145-5p* expression is correlated with calcification of AV tissues explanted from patients undergoing SAVR due to severe AVS and was found to have predictive value for calcification in AVS. Based on the clinical findings of our study, we have further described and explored the functional role of *miR-145-5p* in valvular cells in clinical and in pathological murine models. Of note, the level of *miR-145-5p* was found to be higher in VICs isolated from aortic valve tissue explants from aortic stenosis patients/AVS mice compared to control patients/sham-operated mice and was released into the circulation in a vesicle-associated form. Mechanistically, *miR-145-5p* interferes with the translation of *ZEB2* mRNA by binding to its 3’-UTR. Since it is a negative regulator of the *ALPL* gene, reduction of ZEB2 upregulates *ALPL* expression, resulting in calcification of VICs. The effects triggered by *miR-145-5p in vitro*, such as decreasing viability and migration, increasing inflammation, calcification and simultaneously inducing apoptosis are regarded as prerequisites for the initiation of AVS pathogenesis.

EV-mediated cell-cell crosstalk is gaining more interest in the scientific community due to its implications for non-invasive diagnostics and therapeutic potential. Circulating EVs can be used as biomarkers of disease, as has been shown in various types of cancer and also in cardiovascular diseases, such as atherosclerosis and vascular calcification^19–24^. EVs can be secreted by a variety of cells relevant to cardiovascular diseases, including ECs, platelets, and immune cells. Higher levels of circulating EVs can be observed in patients with classical cardiovascular risk factors such as smoking, hypertension, *diabetes mellitus* and dyslipidemia, and in patients with coronary artery disease (CAD)^25^. Increased levels of circulating endothelium-derived EVs correlate with higher rates of major adverse cardiovascular and cerebral events (MACCE) in patients with stable CAD^25–28^. Not only EV levels, but also more importantly, their cargo composition can predict and influence cardiovascular events. One of the first studies to demonstrate this showed that higher levels of EV-bound miR-126 and miR-199a inversely correlate with MACCE and revascularization-free survival in a patient cohort with stable CAD^28^. These observations gave rise to several mechanistic studies, demonstrating the beneficial and detrimental effects of EVs and their cargo on EC and smooth muscle cells (SMC) in the course of atherosclerosis. Transferred EV-incorporated miRNAs are involved in endothelial regeneration, SMC phenotype switching, osteoblastic differentiation, and vascular calcification^29^. Under calcifying conditions, EVs can be packaged with miRNAs targeting mRNAs for proteins actively involved in osteoblastic differentiation and calcification by a variety of cell types, including SMCs, ECs, and macrophages, demonstrating their important role in vascular calcification processes^29–33^. Recently, we demonstrated that EV-incorporated *miR-122-5p* post-transcriptionally represses *BCL2*, an anti-apoptotic gene, which is central to cell viability and apoptosis^8^. The levels of circulating EV-bound *miR-122-5p* were found to be an indicator of heart function in patients with low or no LVEF improvement after TAVR^8^. EV-incorporated miRNAs have the potential to bind mRNAs that encode for master regulators of osteoblastic differentiation and calcification such as RUNX2 (*Runt-related transcription factor 2*) and ALPL (*Alkaline-phosphatase*). However, the involvement of EV-bound miRNA cargoes in valvular calcification remains unknown.

*MiR-145-5p* belongs to the miR-143/145 cluster of miRNAs, which was reported to be dysregulated (either as a cluster, or one of its members) during essential hypertension, atherosclerosis, CAD, and pulmonary arterial hypertension (PAH)^34–36^. Patients with stable CAD show lower levels of circulating *miR-145-5p* than healthy controls, but patients with unstable angina display elevated levels and *miR-145-5p* levels correlate with infarct size during myocardial infarction. Furthermore, *miR-145-5p* can be transferred between endothelial and vascular SMCs via EVs^34^. Mechanistically, physiological laminar flow induces *miR-145-5p* expression in ECs in a KLF2-dependent manner, which leads to packaging into sEVs and transfers to vascular SMCs, where *miR-145-5p* induces an atheroprotective vascular SMC phenotype^34^. During myocardial infarction, *miR-145-5p* also seems to exert a protective effect by inhibiting apoptosis via the Akt3/mTOR signaling pathway^37–39^. In contrast, the miR-143/145 cluster was found to be upregulated in symptomatic atherosclerotic carotid plaques, when compared to asymptomatic controls, suggesting a role in plaque destabilization^39^. On the other hand, *miR-145-5p* overexpression in ApoE^-/-^ mice via lentiviral vectors was able to reduce aortic atherosclerotic plaque size. These stabilized plaques displayed an increased fibrous cap, more collagen content, and fewer pro-inflammatory macrophages than the plaques from untreated littermates^39^. In patients with PAH, miR-143 and miR-145 levels were found to be higher in pulmonary arterial SMCs (PASMCs) than from healthy controls. *In vitro* experiments demonstrated that *miR-145-5p* influenced PASMC migration and apoptosis, while *miR-145-5p* knockdown ameliorated the development of PAH in a hypoxia induced PAH mouse model *in* vivo^40–41^. Recently, miR-143 was shown to promote valvular calcification by inhibiting matrix Gla protein, (MGP), which itself inhibits calcification and is necessary for valve homeostasis^20^. Thus, the miR-143/145 cluster appears to have multiple roles in the cardiovascular system, ranging from protective to harmful.

*MiR-145-5p* had not previously been investigated in the pathology of AVS. To further validate the involvement of this miRNA in AVS, we examined the AV of mice that had undergone wire-induced injury of the AVs and developed severe AVS. In line with the patient data, we observed an upregulation of *miR-145-5p* in the AV tissue of diseased mice. Since our initial screenings were performed on whole human tissue samples, we sought to identify the cell type that might be responsible for this increase. MiRNA analysis of primary cells from human tissue explants demonstrated a significantly higher expression of *miR-145-5p* in VICs than VECs, suggesting a role during calcification and osteoblastic differentiation. To investigate the function of *miR-145-5p* in VICs, we analyzed its expression *in vitro* under calcifying conditions. A so-called osteogenic medium induces calcification and osteoblastic differentiation of VICs in an ALPL-dependent manner. In our *in vitro* experiments, *miR-145-5p* as well as ALPL levels were significantly upregulated under calcifying stimuli. *In silico* target prediction revealed a specific *miR-145-5p* binding site in the 3’UTR region of ZEB2/SIP1. Of note, the binding of *miR-145-5p* to this binding site in ZEB2 has already been confirmed by a luciferase promotor assay in another experimental setting^42^. Importantly, ZEB2 has been shown to be a transcriptional repressor of *ALPL*^42^. The qRT-PCR analysis confirmed that ZEB2 expression is decreased under calcifying conditions and negatively correlates with *miR-145-5p* and ALPL mRNA levels *in vitro*. To further elucidate the *miR-145-5p*/ZEB2/ALPL axis, we overexpressed *miR-145-5p* in VICs, which led to a significant downregulation of ZEB2 and a consecutive upregulation of ALPL. Furthermore, cells that were simultaneously treated with OM and *miR-145-5p* mimic showed an increased ALPL expression, while *miR-145-5p* ablation diminished ALPL in OM incubated cells, thus indicating an important role of *miR-145-5p* in the modulation of ALPL-dependent calcification.

Since we observed higher levels of *miR-145-5p* in AV tissue- and blood-derived large EVs (IEVs) from AVS patients, we sought to further investigate the transfer of *miR-145-5p* between VICs via EVs. Fluorescence microscopy and copy number experiments confirmed the uptake of PKH26-labelled EVs (a lipophilic membrane dye), fluorescently *labeled-miR-145-5p*, and unlabeled *miR-145-5p* containing EVs by target cells, demonstrating intercellular communication between VICs via EV-bound *miR-145-5p*. The transfer of miRNA cargo between valvular cells might therefore play an important role in AVS initiation and progression. The herein identified regulatory *miR-145-5p-ZEB2-ALPL* axis presents a potential target for the development of new, RNA-based therapies. Furthermore, *in vivo* experiments utilizing EV-incorporated *miR-145-5p* mimics and inhibitors will lead to a better understanding of the involved mechanisms and suggest possible therapeutic strategies. Taken altogether, in this study, using unbiased miRNA profiling, we have identified significantly dysregulated miRNAs in AV tissue from patients with AVS. Among several miRNAs that are commonly dysregulated in mice and humans under AVS, *miR-145-5p* levels were most significantly upregulated and selected for further validation in a larger patient cohort, which subsequently confirmed its expression signature in AVS patients. Interestingly, *miR-145-5p* levels in tissue-derived and circulating large EVs isolated from AVS patients were also higher when compared to control (no AVS) patients. By RNA-sequencing, high-throughput TF array, and proteome arrays utilizing our *in vitro* calcification model of VICs, we revealed that *miR-145-5p* regulates a key process in calcification, i.e. inhibition of ZEB2, a DNA-binding transcription factor that regulates transcription and translation of the ALPL protein, regarded as a hallmark of calcification in valvular and vascular calcification. However, these conclusions from *in vitro* experiments are based on transient overexpression or inhibition of *miR-145-5p* and therefore do not allow us to firmly conclude that *miR-145-5p* is an indispensable factor to promote direct calcification processes in the AV. Importantly, *miR-145-5p* is highly overexpressed in mice and human calcified AV tissues and silencing of *miR-145-5p* reduces apoptosis of VICs and promotes migration and viability of VICs, suggesting that the potential therapeutic effects of pharmacological *miR-145-5p* inhibition for AVS treatment may be transferable into human AVS patients.

## Nonstandard Abbreviations and Acronyms

AVS: aortic valve stenosis
CVD: cardiovascular disease
ECs: endothelial cells
EVs: extracellular vesicles
VICs: vulvular interstial cells
VECs: vulvular endothelial cells
HCAEC: human coronary artery endothelial cell
LV: left ventricle
LVEF: left ventricular ejection fraction
miR: microRNA
miRNA: microRNA
NGS: next generation sequencing
RBP: RNA-binding protein
RIP: RNA immunoprecipitation
SAVR: surgical aortic valve replacement
TAVR: transcatheter aortic valve replacement
TEM: transmission electron microscopy

## Acknowledgements

We thank Ms. Anna Flender and Ms. Sarah Arahouan for their excellent technical assistance.

## Author Contributions

MRH and PRG conceptualized the study. MRH and PRG prepared the manuscript. MRH, PRG, DC, DG, DN, and KB performed experiments. KWJ, JBM, and SU provided scientific input and provided materials. SZ, FB, provided facilities for the murine model and patient-tissue samples. AP, FJ, EL, and GN contributed to the funding of the project, and provided input on the project. All authors have read and approved the final manuscript.

