## Supplementary material for "Aortic valve disease augments vesicular microRNA-145-5p to regulate the calcification of valvular interstitial cells via cellular crosstalk": Online Data Supplement

**Human aortic valve and blood samples**

Calcified, stenotic aortic valve samples were obtained from patients undergoing aortic valve replacement at the University Hospital Bonn due to severe aortic valve stenosis. Non-calcified aortic valve samples were obtained from patients with aortic valve insufficiency undergoing aortic valve replacement surgery, due to aortic root or ascending aorta dilatation. These valves were classified as non-calcified when they did not show any macroscopic signs of calcification or thickening of the cups. Aortic valve tissue samples were stored in 1x PBS on ice immediately after excision for cell isolation and histology and transferred to the laboratory rapidly. To avoid RNA and miRNA degradation, aortic valves designated for RNA isolation were immediately frozen in RNAlater solution (#R0901, Sigma-Aldrich, St. Louis, MO, USA) in liquid nitrogen after excision. Patients with infective endocarditis or post-endocarditic degenerative lesions were excluded as well as patients with bicuspid aortic valves. Further exclusion criteria were: left ventricular ejection fraction below 45%, chronic kidney disease with glomerular filtration rate (GFR) of <60, relevant pulmonary hypertension, any infectious disease, and history of cancer. Informed consent was obtained from all patients. The study protocol was approved by the ethics committee of the University Hospital Bonn (approval number AZ 078/17). Venous blood was obtained prior to surgery and cardiopulmonary bypass and transported to the laboratory on ice.

#### Transthoracic echocardiography

Transthoracic echocardiographic examination was performed on all patients. Biplane-modified Simpson's left ventricular ejection fraction (LVEF) was measured using standard recommendations for echocardiographic quantification. Evaluation of all valves was performed with 2D echo and pulsed- and continuous-wave color Doppler. Peak and mean gradients were calculated using the modified Bernoulli equation, according to the recommendations of the American Society of Echocardiography, using a commercially available ultrasound scanner (Vivid 7, General Electric Medical Health, Waukesha, Wisconsin, USA) with a 2.5 MHz phased-array transducer.

**Isolation of human aortic valve endothelial- and interstitial cells**

Human aortic valve interstitial cells (patVIC) were isolated from heart valves obtained from patients undergoing aortic valve replacement at the University Hospital Bonn. Briefly, valves were washed with 1x PBS, followed by a digestion step using 600 U/ml collagenase II in Endothelial cell growth medium 2 (EBM-2, #CC-3156, Lonza, Basel, CH) for 10 minutes at 37°C. A gentle swab scraping technique led to the removal of endothelial cells, which were collected in a tube and subsequently washed twice with EBM-2. To specifically isolate human valvular endothelial cells (patVEC) positive MACS separation, using human CD105 MicroBeads (#130-051-201,

Miltenyi Biotec, Bergisch Gladbach, D) was performed according to manufacturers' protocol. PatVECs were seeded in EBM-2 and cultured under standard cell culture conditions (37 °C, 5% $CO_2$). Interstitial cells were solubilized by digesting tissue pieces in 10 ml collagenase II (600 U/ml) in a basic patVIC medium (DMEM/GlutaMAX, #61965026, Gibco, Carlsbad, CA, USA, supplemented with 10% FBS, 1% Penicillin/Streptomycin and 50μM sodium bicarbonate) for 12 h (37 °C, 5% $CO_2$) under gentle rotation. Samples were filtered through a cell strainer (100 μm pore size), and cells were pelleted and washed twice with patVIC medium. The isolated patVICs were cultured in patVIC medium under standard cell culture conditions (37 °C, 5% $CO_2$). Cells in low passages were used for experiments.

**Culture and characterization of valvular cells**

VICs and VECs were purchased from Lonza (Basel, CH). VICs were cultured under standard cell culture conditions (37 °C, 5% CO2) in DMEM/GlutaMax, including 10% fetal bovine serum (FBS, #A3160802, Gibco) and 1% Penicillin/Streptomycin (PS). VECs were cultured in EBM-2, including the supplement kit: EGM-2 MV (#CC-3202, Lonza). Cells were passaged using the Lonza Clonetics ReagentPack (#CC-5034 Lonza) and used for experiments in passages 3-6. For experiments VICs and VECs were seeded in 12- (VICs: 25.000 cells/well; VECs: 50.000 cells/well) and 6-well-plates (VICs: 60.000 cells/wells).

In order to subject patVICs, patVECs, VICs, and VECs to immunofluorescence (IF) analysis the cells were seeded on coverslips and fixed with 4% Paraformaldehyde for 30 minutes followed by three washing steps with 1x PBS and 10 minutes permeabilization with 0.25% Triton-X. After three further washing steps, coverslips were blocked using 1% BSA-Glycin in PBS containing 0.1% Tween-20. Cells were incubated for two hours with primary antibodies (α-SMA, dil.: 1:100, #ab7817; Von Willebrand Factor, dil.: 1:100, #ab6994; Vimentin, dil.: 1:1000, #ab45939, VE-cadherin, dil.: 1:500, #ab33168; all Abcam Cambridge, UK) diluted in 1% BSA-Glycin in 1x PBS containing 0.1% Tween-20. Subsequently, the primary antibodies were removed by three rinses with PBS in preparation for incubation with the secondary antibody (#115-165-146, #111-225-144, #712-165-153, Dianova, Hamburg, D) for one hour. Afterward, coverslips were washed again with PBS, before cells were counterstained and embedded with Vectashield mounting medium with DAPI (#H-1200-10, Vector laboratories, Burlingame, USA). Observations were performed with fluorescence microscopy (Axio Observer Inverted Microscope, Zeiss, Jena, D). Furthermore, interstitial and endothelial cell markers were quantified using quantitative reverse transcription-polymerase chain reaction (qRT-PCR).

**Osteogenic differentiation of valvular interstitial cells**

Osteogenic differentiation was induced by treatment with osteogenic medium OM (DMEM+GlutaMax, 1% FBS, 1% Penicillin/Streptomycin; 10 nmol/L beta-glycerolphosphate, 50 µmol/L ascorbic acid, 10 µmol/L dexamethasone) for 7 days. Normal cell growth medium was used as control (CTM; DMEM+GlutaMax, 1%FBS, 1% Pen-Strep).

**Transfection**

*MiR-145-5p* miRCURY-LNA inhibitor (#YI04102423 Qiagen, Hilden, Germany) and *miR-145-5p* miScript mimic (#MSY0000437, Qiagen) were used for transfection. Transfection was performed on 12-well plates using 2µl/well Lipofectamine RNAiMax (#13778150, ThermoFisher Scientific, Waltham, MA, USA) and oligonucleotides according to the manufacturer's instructions. Different concentrations of oligonucleotides were tested, and the statistically significant results were used for further experiments. Before transfection, cells were washed with 1x PBS, and the medium was changed to DMEM containing 10% FBS without antibiotics. Lipofectamine and oligonucleotides were diluted in Opti-Mem (#11520386, Gibco) and after incubation added to the cells. The transfection medium was removed after 24h and cells were washed with 1x PBS. Cells were either kept on normal growth medium for another 24h and were subsequently harvested or were treated with normal growth medium or OM for indicated time points.

**EV isolation from valvular tissues and cells**

Aortic valve tissue-derived EVs were isolated as follows: AV tissue was dissected into small parts under sterile conditions. A 6-well plate was prepared with 2mL DMEM (#11039-021, Gibco) per well, the snippets were distributed equally into the wells and were incubated for overnight under gentle mixing on a rotator at 37° C, 5% $CO_2$. EV isolation from the supernatant was performed by differential (ultra-) centrifugation following the protocol in the supplement. Pellets after 20.000g and 100.000g centrifugation steps containing large and small EVs, respectively, were resuspended in 25 µl 1xPBS and used for further experiments and characterization of EVs.

For cell culture-derived EVs, cells were incubated in a cell culture medium containing exosome-depleted FBS for 24h before isolation. The supernatant was used in further centrifugation steps according to our protocol. EVs were characterized by nano tracking analysis (NTA), western blot (WB), and transmission electron microscopy (TEM) in accordance with ISEV guidelines.

EVs from patient blood were isolated as follows: Citrate blood (S-Monovette, Sarstedt, D) was collected from patients with and without aortic valve stenosis and transported to the laboratory on ice. Platelet-depleted plasma was generated by three centrifugation steps (2 x 2500 g, 15 min, 4 °C; 1 x 3000 g, 15 min, 4° C). Large EVs were isolated from platelet-depleted plasma by centrifugation at 20.000 g. The supernatant was washed at

100.000g and small EVs were pelleted by ultracentrifugation at 100.000 g.

**Vesicle–RNA degradation assay**

Large EVs were resuspended in PBS. The sample from the untreated group was left on ice as a control. To digest protein, 45 µl Proteinase K (Thermo Fisher Scientific, #25530049) was added to one sample. Additionally, 25 µl Triton X-100 (Sigma-Aldrich, #T8787) was added to another sample for 30 minutes at 37°C to disrupt the membrane bilayer of the large EVs. Afterwards, all samples were treated with 5 µl RNase A (Thermo Fisher Scientific, #AM2271) for 10 minutes at 37°C. Finally, the samples were lysed with Qiazol, and RNA was isolated for qRT-PCR analysis (normalized to spiked-in cel-miR-39).

**TaqMan miR array**

MicroRNA arrays were performed using the TaqMan Array Human MicroRNA Card A v2.0 (Applied Biosystems #4398965) in a 7900 HT fast real-time PCR instrument (Applied Biosystems). Total RNA (500 ng) from the tissue of patients with AVS (n=4) and Controls (n=4) was converted into cDNA by use of the TaqMan MicroRNA Reverse Transcription Kit and specific Megaplex RT Primers (Applied Biosystems, #4399966). Subsequently, preamplification was performed using 2.5 µL of the RT product, Megaplex PreAmp Primers (Applied Biosystems, #4399233), and the TaqMan PreAmp Master Mix (Applied Biosystems, #4488593). 9 µl of the preamplification product mixture was used for the final amplification with the MicroRNA Array Card and TaqMan Universal Master Mix II. The amplification curves were analyzed with the program RQ manager Version 1.2.1 (Applied Biosystems) to calculate CT values, using the same threshold across all respective plasma samples. Only miRs that were stably expressed in all three samples were included in the final analysis. ΔΔCT values were calculated by normalizing to *RNU6B,* which was the most stably expressed endogenous control RNA across the plasma samples.

**RNA isolation and qRT-PCR**

RNA of cultured cells and tissue samples were isolated using a TRIzol (#15596026 ThermoFisher Scientific) and phenol-based protocol. After purification RNA was diluted in DNAse/RNAse free water and RNA concentration was measured via NanoDrop (ThermoFisher Scientific). For EV-bound RNA, isolation was performed using the miRNeasy Mini Kit (#217004, Qiagen) according to the manufacturers' instructions. RNA was converted into cDNA using Omniscript RT Kit (#205111, Qiagen) for gene expression analysis, for microRNA we used TaqMan microRNA Reverse Transcription Kit (#4366596, ThermoFisher Scientific) and microRNA-specific primers. A quantitative polymerase chain reaction was carried out on a 7900HT thermocycler (Applied Biosystems) with with the following condition: 95°C for 10 min followed by 40 cycles of 95°C for 15 sec and 60°C for 1 min. cDNA was diluted to 1ng/µl and 9µl were pipetted per well. 1µl primer was mixed with another 10µl of Universal MasterMix II (#44-400-49 ThermoFisher Scientific) or Gene Expression MasterMix (#43-700-74, ThermoFisher Scientific). CT values above 40 cycles were defined as undetectable. Data were analyzed using the ddCT method.

**MiR target prediction**

The miRNA database and target prediction tool TargetScan (http://www.targetscan.org) was screened to identify potential miRNA targets involved in alkaline phosphatase regulation.

**Luciferase reporter array**

For analysis of transcription factor activity, a Cignal Finder Reporter Array (Qiagen, #336821) was used as recommended by the manufacturer. VICs were transfected with siRNAs as described previously. After transfection, 40.000 VICs/well were seeded into 96 well plates from Qiagen. Cells were seeded in 50µl OptiMEM containing 10% FCS while each well contained 100µl OptiMEM supplemented with Lipofectamine RNAiMAX. Cell culture plates were incubated for 4h (humidified atmosphere, 5% $CO_2$, 20% $O_2$, 37°C). Subsequently, the medium was changed to a VIC culture medium as previously described and culture plates were incubated for another 24 h following transfection in the culture plates. Luciferase signals were measured with a GloMax – Multi Detection System from Promega.

**Alizarin Red staining**

*In vitro* calcification was assessed via Alizarin Red S Staining (#A5533, Sigma-Aldrich). Cells were fixed with 4% PFA for 15 minutes at 4°C. Cells were subsequently washed with 1x PBS. For calcification staining, cells were incubated in 0.02 mg/L Alizarin Red S staining solution for 45 minutes, washed, and light microscopy with a ZEISS Axiovert 200M microscope was performed to visualize calcium-phosphate deposition.

**Aortic stenosis model and Echocardiography**

10–12-week-old male C57BL/6-J (wild-type) mice were purchased from Janvier Labs, France. All animal experiments were performed according to institutional guidelines and the German animal protection law, and the operations were performed according to a protocol that was published previously[42-43]. Fujifilm VisualSonics Vevo 2100 Ultra High-Frequency Imaging Platform was used for the assessment of the functional analysis. Mice were anesthetized with 1.5% isoflurane, with continuous monitoring of the electrocardiogram, respiratory rate, and body temperature. Standard analyses of aortic valve peak velocity and ejection fraction were performed.

The induction of aortic valve stenosis in mice, briefly described, was achieved blindedly by inserting a coronary wire into the left ventricle (LV) via the carotid artery, which was then pushed/pulled through the valve and rotated repeatedly to induce AVS (Figure S2A). In detail, the mice were anesthetized by intraperitoneal injection of 150 mg/kg ketamine and 16 mg/kg xylazine. The right carotid artery was exposed by blunt dissection and the blood flow was stopped using ligature loops. For mild and moderate injury, a straight guide wire with a shortened and soldered tip (Abbott HI-TORQUE 0.014) was used. For severe injury, a conventional guide wire with a 15° angled tip (ASAHI INTECC MIRACLEbros 6) was used. The wire was introduced into the left ventricle under echocardiographic guidance, passed through the aortic valve, advanced into the left-ventricular apex, and pulled back into the left-ventricular outflow tract, just below the level of the aortic valve. This resulted in an amplitude of 4–5 mm. Hereafter, the wire was rotated within the valve with a speed of two rotations per second. After the injury, acute aortic valve regurgitation was immediately assessed using a color-Doppler ultrasound. The wire was removed and the carotid artery was ligated. The sham procedure was performed in the same fashion, but the wire was only inserted into the right carotid artery and not advanced across the aortic valve into the left ventricle.

Aortic valve peak velocity was measured in the suprasternal view with a pulse-wave-Doppler, using an angle correction between 40° and 50°. An increase in peak velocity of 15–50% from baseline was defined as mild aortic stenosis, 50–75% as moderate, and > 75% as severe aortic valve stenosis.

Left ventricular ejection fraction, fractional shortening, and ventricular volumes were measured in parasternal long-axis views using the Vevo LV-Trace function. Wall thickness was measured in parasternal long- and short-axis M-Modes. Aortic valve regurgitation was imaged using color-Doppler mode in the parasternal long-axis and suprasternal views.

**Nanoparticle tracking analysis**

Size and concentration distribution of EVs from the plasma and endothelial cells were performed by using nanoparticle-tracking analysis (NTA) with a Nanosight NS 300 (Malvern Instruments, UK). Each sample was recorded five times for 60 seconds at a speed level of 20. The analysis was performed by setting a detection threshold of six. PBS was used to perform a background measurement, in order to confirm the absence of residual particles. The NTA software (version 3.1 Build 3.1.46) was used to record and analyze the samples.

**Electron microscopy**

For transmission electron microscopy (TEM), large EVs from plasma or CMs were pelleted by centrifugation (20,000 xg, fixed in 1.25% glutaraldehyde in 0.1 M cacodylate buffer overnight, dehydrated with ethanol and propylene oxide, and embedded in Epon 812 (Serva), also overnight. After double-contrast staining with uranyl acetate and an aqueous lead solution, images were taken with a CM 10 electron microscope (Philips).

**EV-incorporated *miR-145-5p* uptake by vulvular cells**

VICs were transfected with green488-labeled *miR-145-5p* (#L-00211, RiboLink, Riboxx) using Lipofectamine RNAiMax. The day after transfection, VICs were washed with sterile 1x PBS, and the medium was switched to a normal medium containing 10% exosome-depleted FBS (#2720801, Gibco). After incubation for 24h, the supernatant was used to generate large EVs following our isolation protocol. Target VICs were incubated with these lEVs for 6h. After incubation, cells were fixed with 4% PFA for 15 minutes (4°C) and nuclei were counterstained with DAPI (Vector laboratories). In the second set of experiments large EVs, generated from untransfected VICs, were labeled with PKH26 (#MIDI26, Sigma-Aldrich,) according to the manufacturer's instructions. PKH26-labeled EVs were washed twice with 1x PBS. To evaluate the uptake of EVs into cultured cells, VICs were incubated with PKH26-labeled EVs for 6 hours. After incubation, cells were fixed with 4% PFA for 15 minutes (4°C) and nuclei were counterstained with DAPI (Vector laboratories). A Zeiss Axiovert 200M microscope and AxioVision software was used to visualize the uptake of EV-incorporated *miR-145-5p* or PKH26-labelled EVs into recipient cells.

**Copy Number analysis of *miR-145-5p* and uptake into target VICs**

After 24h of transfection of donor VICs with *miR-145-5p* mimic or mimic negative control (Qiagen), the medium was switched to a normal medium containing 10% exosome depleted FBS (Gibco) for 24h. Large EVs were isolated from the medium and re-dissolved in sterile PBS. A small fraction of EVs was used for RNA isolation while the major rest was incubated with recipient VICs for 24h. The absolute expression of *miR-145-5p* was determined by obtaining standard curves with different concentrations of *miR-145-5p* mimic as templates. Logarithmic values of oligonucleotide concentrations and CT values were plotted. The copy number was calculated from the oligonucleotide concentration and the molecular mass of the transcript.

**Transwell co-culture assay**

VECs were transfected as described above, before performing the co-culture assay. Once the cells were confluent, *miR-145-5p* mimic-, inhibitor- and control-transfected VECs were seeded in the co-culture insert. VICs were seeded in 24-well plates. On the day of the co-culture experiment, the medium in the co-culture insert was exchanged for a serum-free basal medium to induce the generation of large EVs from the VECs in the insert. Inserts with VECs were then placed in the plate wells containing VICs and both types of cells were then in a co-culture system via the porous transwell membrane (1 µm pore size and incubated for 24 hours. VICs from the 24-well plates were then collected in TRIzol for RNA isolation and subsequent qRT-PCR analysis.

**MTT-based cell-viability assay**

30000 cells were seeded overnight per well in a 6-well plate and transfected with the respective siRNAs for 48 hours, followed by isolation of large EVs to incubate with recipient cardiomyocytes for 24 hours. A total of 30,000 recipient VICs were transferred to the 96-well flat bottom plate before incubation overnight at 37°C, 5% $CO_2$, in an incubator. The cells were incubated with 10 µL MTT solution (3-(4,5-dimethylthiazol-2-yl)-2,5-diphenyltetrazolium bromide), as designated in the MTT Cell Growth Assay Kit (Millipore, #CT01), in culture medium for 4 hours at 37°C. Then, a formazan formation reaction was performed and changed colorimetrically by the addition of 100 µL isopropanol with 0.04 N HCl to each well, followed by gentle tapping to mix the solution. The absorbance was measured on an ELISA plate reader with a test wavelength of 570 nm and a reference wavelength of 630 nm.

**Caspase 3/7 activation assay**

Caspase-3/7 activity in human VICs was assessed using the Apo-ONE Homogeneous Caspase-3/7 kit (Promega, # G7792), according to the manufacturer's instructions, 48 hours after transfection. Briefly, the caspase substrate Z-DEVD-R110 was diluted 1:100 in Apo-ONE Homogenous Caspase 3/7 buffer and incubated with the cells for 1 h at 37°C. For induction of apoptosis, VICs were incubated with $H_2O_2$ (in DMSO, 100 µM from a 500 mM stock solution, Sigma-Aldrich, #H1009-100ML) for 6 hours before the addition of the substrate. Fluorescence was measured at a wavelength of 521 nm.

**Human apoptotic protein profiling**

The Proteome Profiler™ Array Human Apoptosis Array Kit (R&D Systems) was used according to the manufacturer's instructions 48h after siRNA treatments to VICs. Firstly, provided arrays were blocked for 1 h with array buffer and then incubated with cell lysates overnight at 4 °C. Arrays were incubated with Detection Antibody Cocktail for 1 h and subsequently with Streptavidin-HRP for 30 min at RT. Chemi Reagent Mix was used for visualization with the Amersham Imager 600 (GE Health care). Band intensities were quantified in ImageQuant TL (GE Healthcare).

**Western blotting**

As a control for cell EVs in immunoblotting, cell lysate was used, whereas, in the case of EVs from cell culture, cell lysate and conditioned media (growth media without growth-media supplements, PromoCell, #C-22020) were used as a control. Lysates were ultrasonicated for 10 min and the protein concentration was assessed with a Qubit-4 Fluorometer (Thermo Fisher Scientific) by use of a Qubit™ Protein Assay Kit (Thermo Fisher Scientific, #Q33211), according to manufacturer's instructions. For quantification of the EV protein markers as well as for the confirmation of the absence of tubulin in EVs, 50 µg of lysates were used and for the confirmation, 30 µg of proteins were diluted 2:1 in 3x Laemmli buffer and loaded onto an SDS-PAGE gel (Biorad, #456-1084 and 456-1024; Mini PROTEAN System, Biorad). Subsequently, the protein was transferred onto a Roti-NC nitrocellulose membrane (Carl Roth GmbH, HP40.1) and blocked with 5% BSA (Sigma-Aldrich) for one hour. Mouse Anti-CD63 antibody (Abcam, #ab59479, RRID:AB_940915) 1:1000, mouse Anti-CD31 antibody (Abcam, # ab28364) 1:1000, anti-ZEB2 antibody (Abcam, #2128)) 1:1000, Anti-human Flotilin (Abcam, #ab10241), 1:1000, ab10241, Anti-tubulin (Cell signaling, #ab10241), 1:1000, and mouse anti-β-Actin antibody (Sigma-Aldrich, # A1978, RRID:AB_476692) 1:2000 in 5% BSA were used to stain the membrane overnight at 4°C. After extensive washing with 0.1% TBST, the membrane was incubated with an HRP-conjugated rat monoclonal anti-mouse-IgG antibody (Sigma-Aldrich, #A9044, RRID: AB_258431) 1:3000 or Anti-Rabbit HRP (Abcam, #4750.1) 1:1000 in 5% BSA for 1 hour at RT. After washing again with 0.1% TBST, the membrane was developed with ECL primer western-blotting detection reagent (Sigma-Aldrich) and imaged by use of a ChemiDoc MP imaging system (Biorad). The images were analyzed by the software ImageJ (NIH, USA).

**RNA-seq data analysis**

FASTQ files were preprocessed with fastp (version 0.21.0) using the default setting. After preprocessing of sequencing reads, STAR (version 020201) was used to map the reads to the reference genome (GRCh38.103). To calculate counts per million (CPM) values and derive differentially expressed genes, the R package, edgeR [3] (version 3.30.3), was used. For the pre-processing of the data we used the nf-core pipeline for RNA-Seq. After adapter and quality trimming (Trim Galore), reads have been aligned to the human genome with STAR. Then, featureCounts was used to quantify gene abundances from the alignment coverage where only uniquely mapped reads were accounted. Gene Ontology and KEGG pathway analyses were performed using DAVID (https://david.ncifcrf.gov/home.jsp)[3] and LSKB software (http://www.lskb.w-fusionus.com)[4] (World Fusion.inc, Tokyo Japan). Venn diagrams were drawn via http://bioinformatics.psb.ugent.be/webtools/Venn/.

**Supplemental Figures with Legends:**

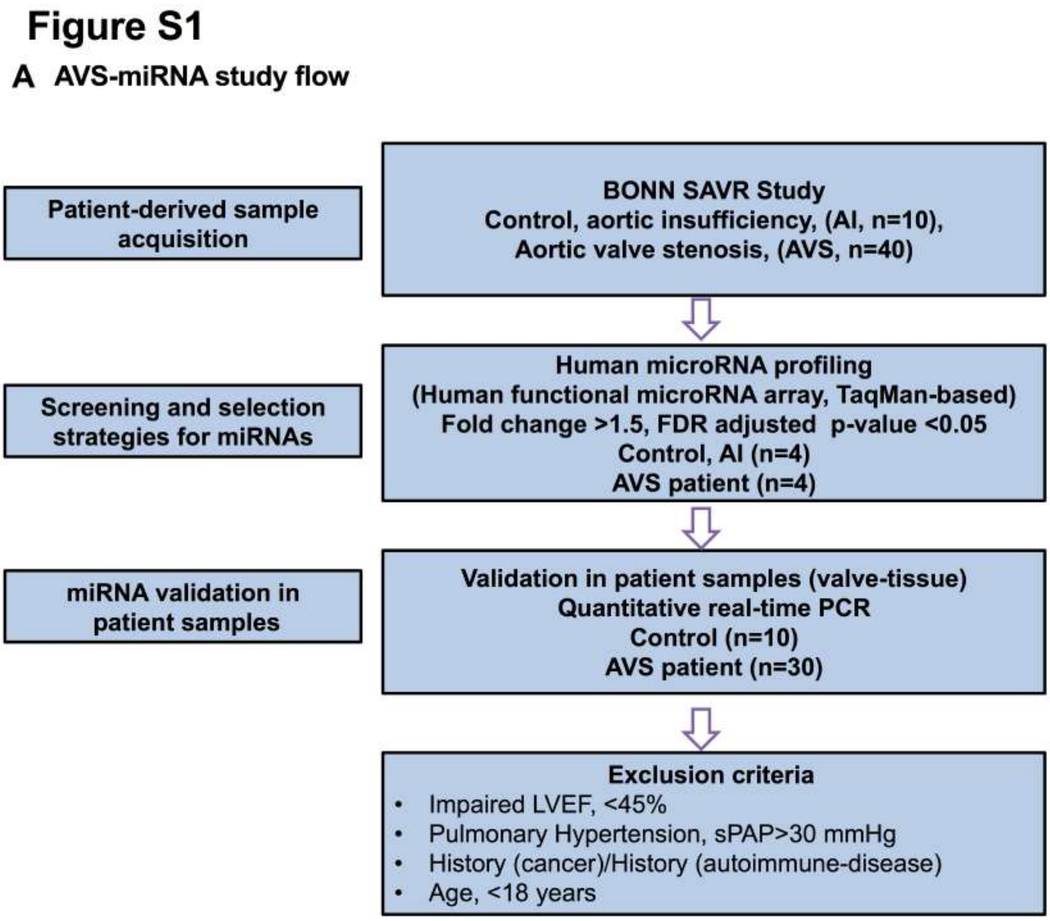


**Figure S1. *miR-145-5p* is associated with LVEF improvement and clinical outcome.**

(A) Schematic representation of the clinical miRNA study. Patient samples were obtained from a well-documented Bonn SAVR biobank. SAVR, surgical aortic valve replacement; FDR, false discovery rate; PCR, Polymerase chain reaction.

#### Figure S2

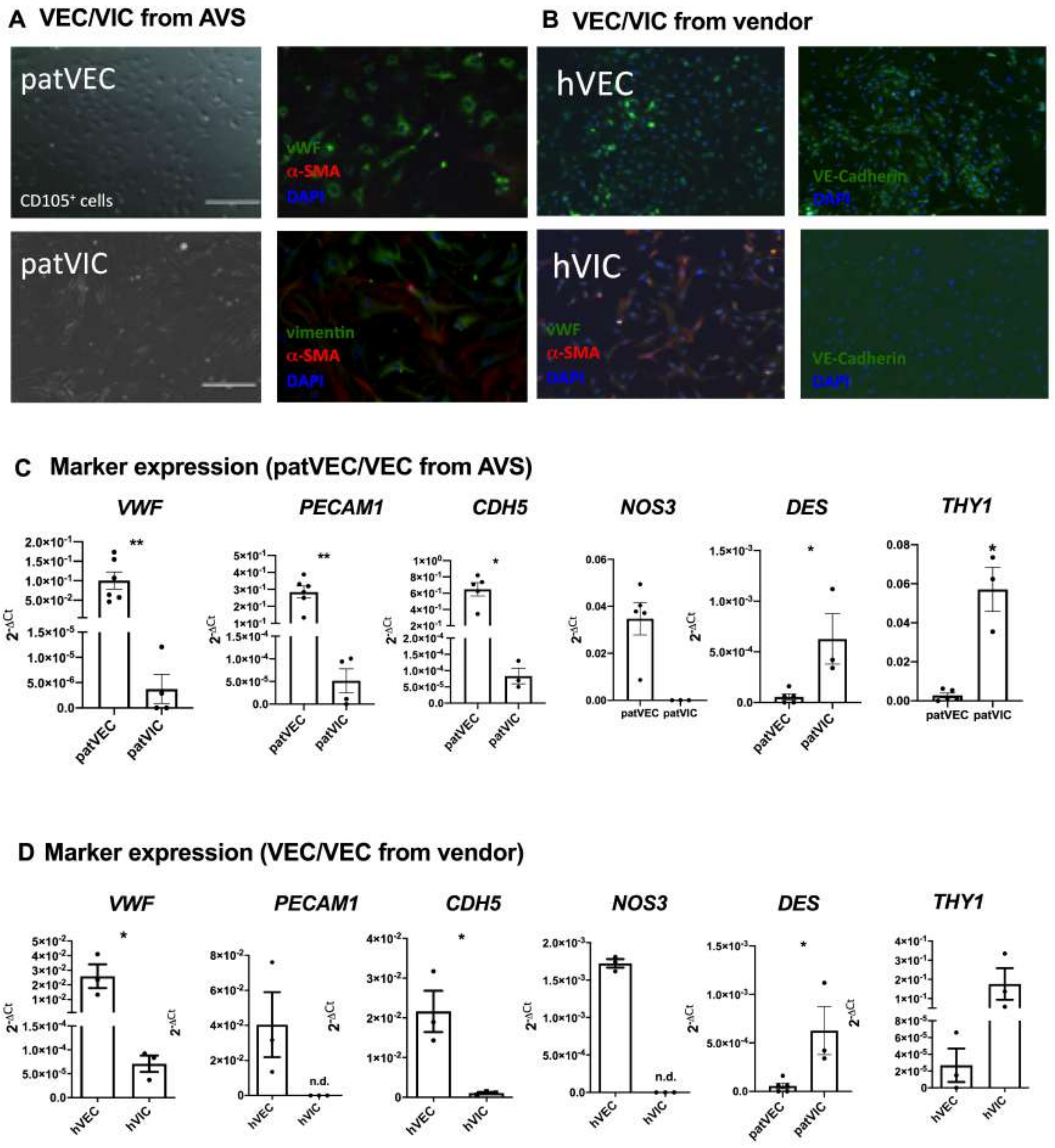


**Figure S2. Comparative analysis of patient-derived and commercially-available valvular cells.**

(A-B) Characterization and comparison of isolated and commercially available aortic valve cells by using different surface markers, corresponding to VECs- and VICs-derived from AVS patients that underwent SAVR and compared with the corresponding controls and purchased cells. Representative immunofluorescence images of isolated VECs (patVECs) and VICs (patVICs) by using their characteristic markers. (C-D) Expression of characteristic markers of isolated valvular cells, VECs and VICs, isolated from patients with AVS post SAVR and commercially-available cells. Data are depicted as Student t-tests. (****$p<0.0001$, **$p<0.01$, n=3-5, two-tailed, unpaired). VICs, valvular interstitial cells; VECs, valvular endothelial cells; SAVR, surgical valve replacement; vWF, von-Willebrand factor; a-SMA, alpha-smooth muscle actin; DAPI, 4′,6-Diamidin-2-phenylindol; VE-Cadherin, vascular endothelial cadherin; PECAM1, platelet endothelial cell adhesion molecule; CDH5, cadherin 5/VE-Cadherin; NOS3, nitric oxide synthase 3/eNOS; DES, desmin; THY1, Thy-1 Cell Surface Antigen/CD90; DES, desmin; THY1, Thy-1 Cell Surface Antigen/CD90.

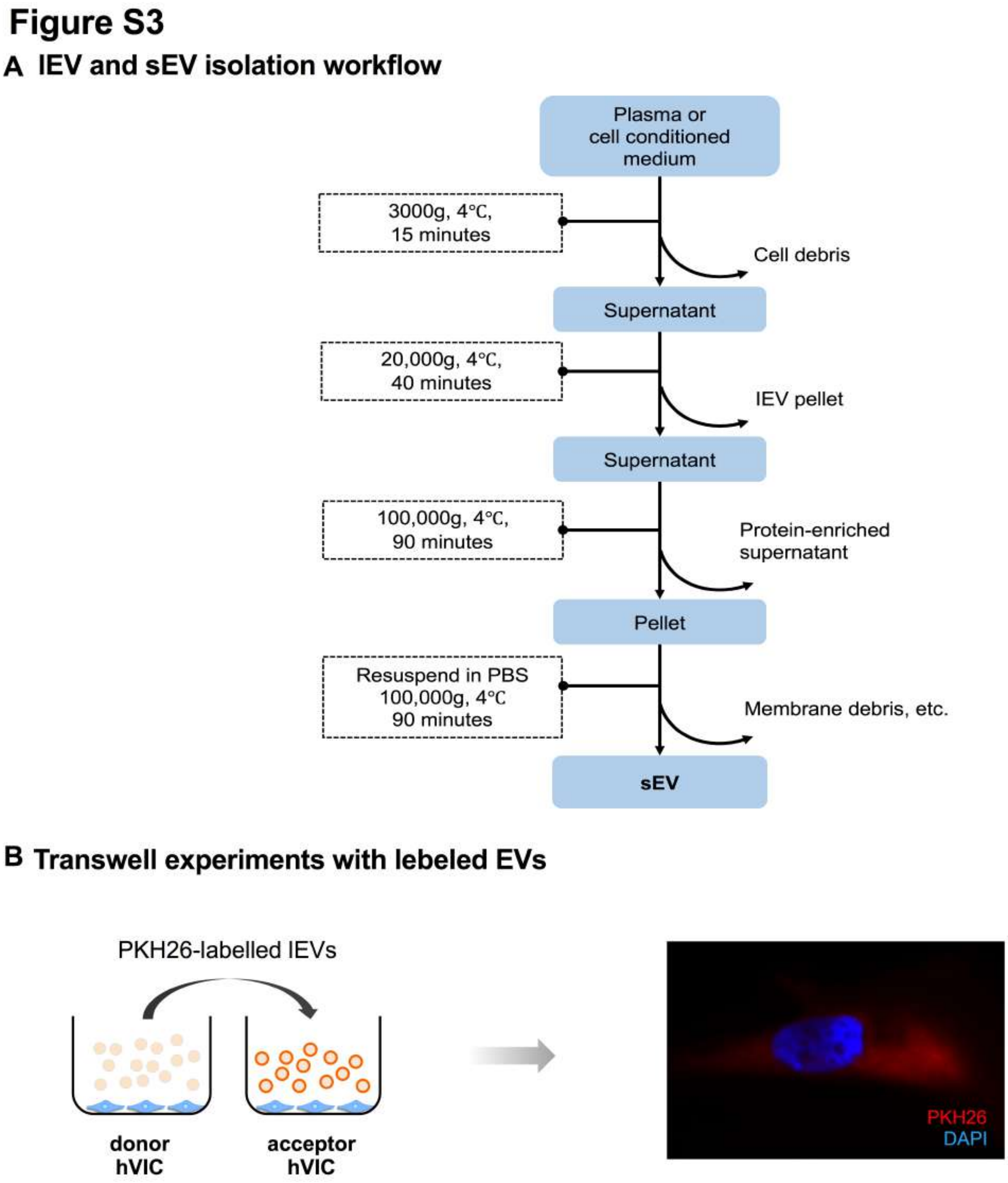


**Figure S3.** ***MiR-145-5p* may shuttle to recipient VICs via vesicular transfer.**

(A) Schematic flows of the isolation of EV population from AV-tissues, plasma, and cell culture. (B) EV incorporation into VICs from the recipient to donor incorporation experiment by VICs. PKH26-labeled large EVs were co-incubated for 24 hours to absorb EV in recipient VICs. EVs, extracellular vesicles; AVS, aortic valve stenosis; lEV, large EV; sEV, small EV; hVICs, human vascular interstitial cells.

### Figure S4

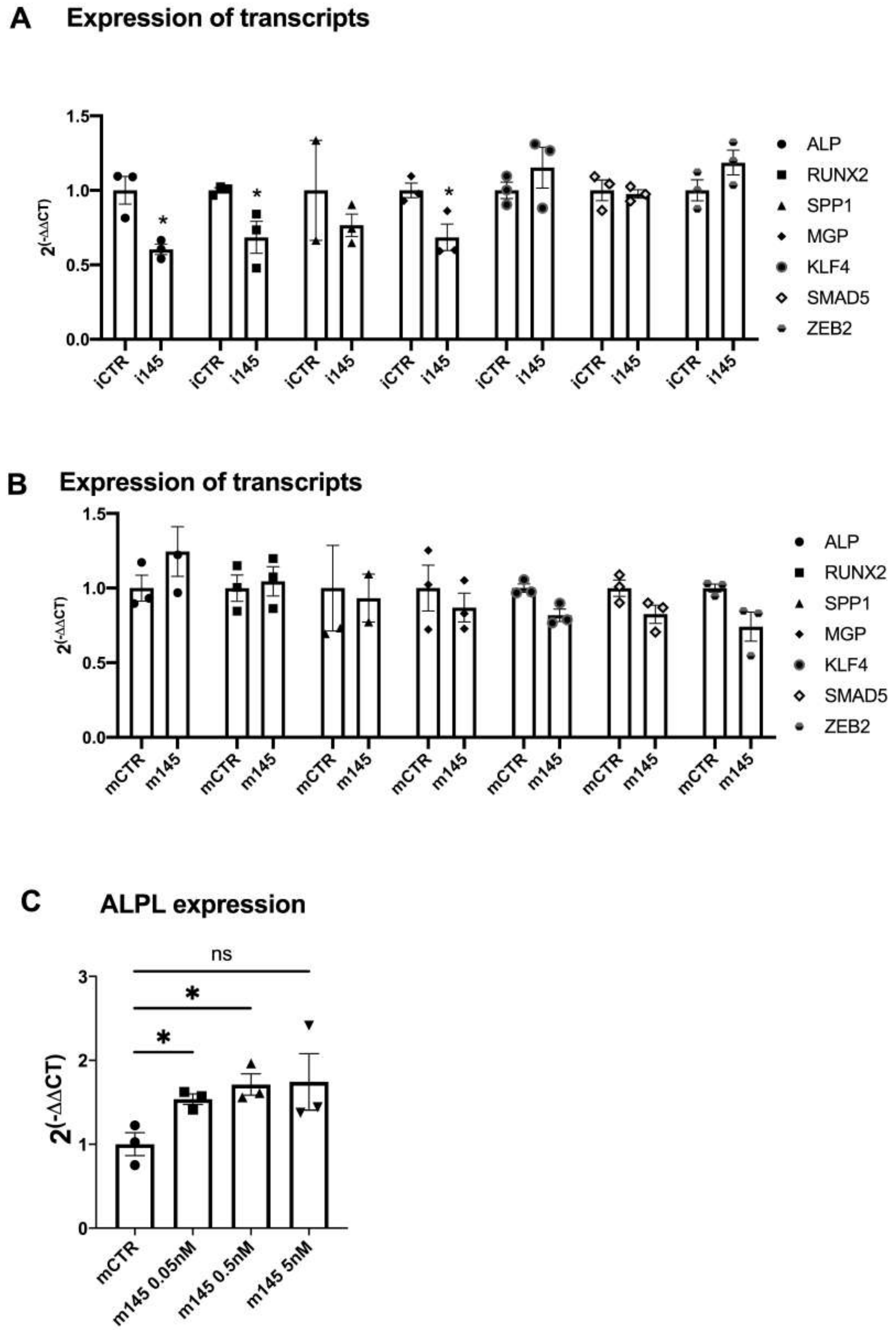


**Figure S4.** ***MiR-145-5p*** **may regulates valvular calcification genes.**

(A-B) *In vitro* calcification experiment in VICs followed by quantification of calcification genes overexpression and knockdown quantification of *miR-145-5p* by small oligonucleotides in VICs. Expression data for calcification genes in VICs have been compared with control and depicted as *$p<0.05$, n=3, by 1-way ANOVA with Bonferroni multiple comparisons test. (C) A Taqman-based quantification of gene expression of ALPL, as a hallmark of calcification of VICs, upon different concentrations of *miR-145-5p* mimic transfection followed by induction of *in vitro* calcification by using osteogenic medium (OM) after 7 days. Expression data for ALPL mRNA in VICs have been compared with control media and depicted as $^{ns}p>0.05$, *$p<0.05$, ***$p<0.001$, n=3, by 1-way ANOVA with Bonferroni multiple comparisons test. mCRT, mimic controls; m145, mimic-145; ALPL, Alkaline Phosphatase; RUNX2, Runt-related transcription factor 2; SPP1, Secreted Phosphoprotein 1; MGP, Matrix Gla Protein; KLF4, Kruppel Like Factor 4; SMAD5, SMAD Family Member 5; and ZEB2, Zinc Finger E-Box Binding Homeobox 2; AVS, aortic valve stenosis.

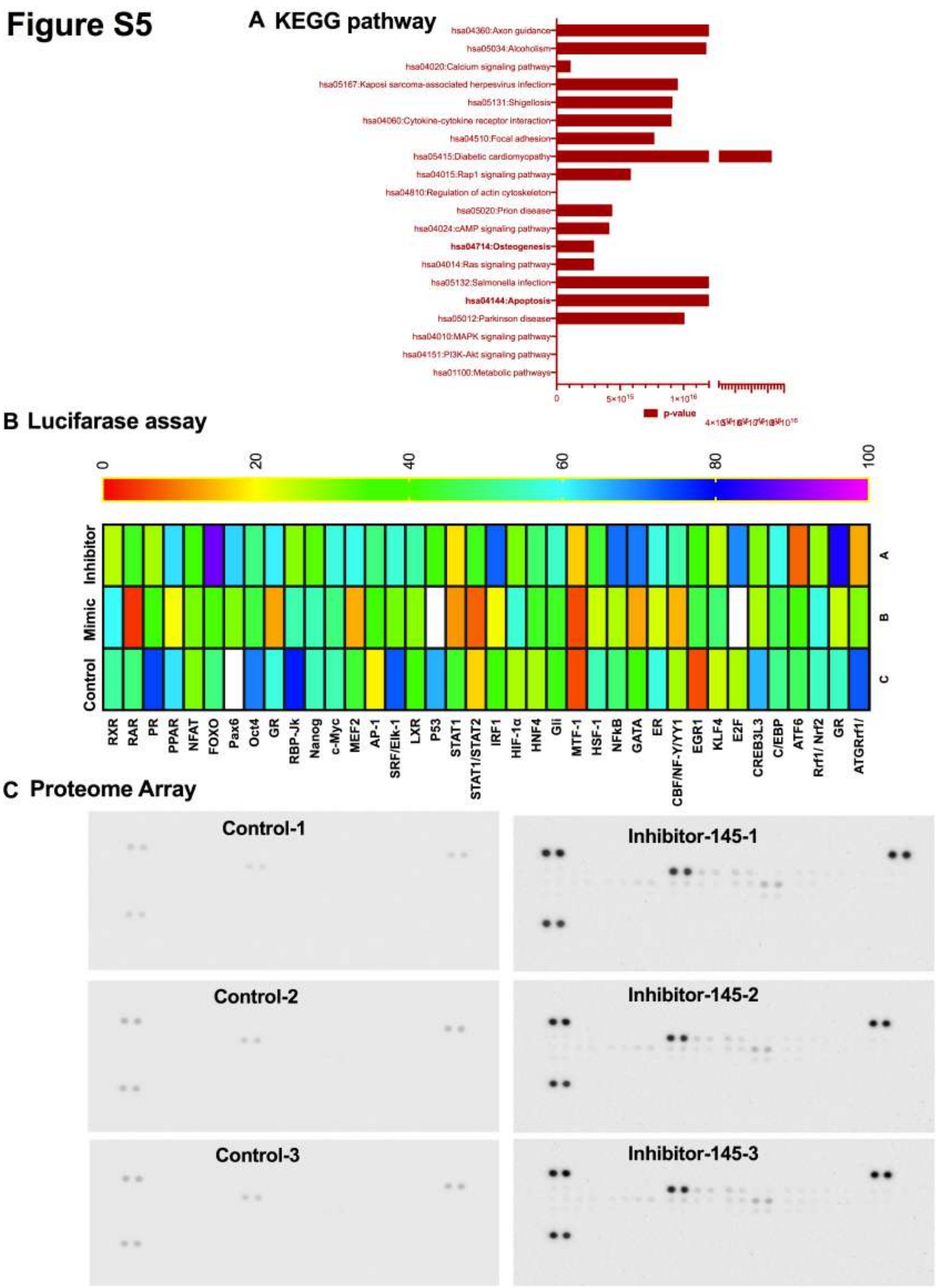


355

**Figure 5. Transcriptomic profiling of calcified VICs supports the regulation of calcification and apoptosis by *miR-145-5p*.**

(A) KEGG pathway analysis with dysregulated genes in RNA sequencing analysis showing that osteogenesis and apoptosis are among the dysregulated pathways. (B) Heatmap with profiling of dysregulated transcription factors by using a high throughput transcription factor activity reporter luciferase assay upon oligonucleotide-mediated depletion or overexpression of *miR-145-5p*/control in VICs was performed 48h after transfection. Depicted are the activities of 45 transcription factors in inhibitor *miR-145-5p* treated cells, mimic *miR-145-5p* and control cells (control-treated) 48h after transfection with the reporter constructs. Pro-Caspase-3, Caspase-3, HSP27, HIF1a, and other apoptosis-related pathways are significantly upregulated. (E) Pro-Caspase-3, Caspase-3, HSP27, HIF1a, and other apoptosis-related pathways are significantly upregulated. Data represent mean ± SEM (****$p<0.0001$, $n\geq6$). ZEB2, Zinc Finger E-Box Binding Homeobox 2; RUNX2, Runt-related transcription factor 2; ALPL, Alkaline Phosphatase; HSP27, Heat shock protein 27; HIF1a, Hypoxia-inducible factor 1-alpha; CTM, control media, OM, osteogenic medium; UTR, untranslated region; VICs, valvular interstitial cells; miRs, microRNAs; AVS, aortic valve stenosis.

**Suplementary Table legends:**

**Table S1. Transcription factor array**

**Table S2. Proteome array**
